# ALTERNATIVE SPLICING OF A CODING GENE PRODUCES A NUCLEAR REGULATORY LONG NON-CODING RNA

**DOI:** 10.64898/2026.03.31.715525

**Authors:** Florencia S. Rodríguez, Pablo Mammi, Federico E. Aballay, Loreana Pulichino, Rocío S. Tognacca, María Florencia Legascue, Nicolás Gaggion, Lucas Servi, Maria Kalyna, Andrea Barta, Federico Ariel, Martín Crespi, Ezequiel Petrillo

**Affiliations:** Instituto de Fisiología, Biología Molecular y Neurociencias, CONICET - Universidad de Buenos Aires, Facultad de Ciencias Exactas y Naturales, Buenos Aires 1428, Argentina; Institute of Plant Science (IPS2), Paris-Saclay University, France; Apolo Biotech, Santa Fe, Argentina; Institute of Molecular Plant Biology, Department of Biotechnology and Food Science, BOKU University, 1190 Vienna, Austria; Max Perutz Labs, Medical University Vienna, Vienna BioCenter (VBC), Vienna, 1030, Austria

## Abstract

Alternative splicing (AS) is traditionally understood to increase transcript and protein diversity by generating multiple coding isoforms from a single gene. In addition, many alternative isoforms are degraded through non-sense mediated decay (NMD). Furthermore, we show here that AS also produces nuclear long non-coding RNAs (lncRNAs) from protein-coding genes, which can serve regulatory functions rather than expanding proteomic complexity. Focusing on the *Arabidopsis thaliana* SR protein gene *At-RS31*, we found that a non-coding isoform (*mRNA3*) accumulates in the nucleus under dark conditions. This transcript binds the protein product of its own transcriptional unit, modulating splicing decisions and balancing gene activity in response to environmental cues. Overexpression of *mRNA3* down-regulates the action of At-RS31 on its target genes *in trans* and restores the phenotype induced by intron-less *At-RS31* accumulation. Further sub-cellular fractionation and RT-PCR analyses show that many SR genes generate nuclear-retained non-coding isoforms, especially under dark conditions, suggesting a widespread mechanism. These findings redefine the role of AS in plants, highlighting its capacity to generate regulatory lncRNAs to fine-tune gene expression beyond protein diversification.

**Highlights:**

- Alternative splicing generates regulatory lncRNAs from protein-coding genes
- A nuclear-retained isoform forms an autoregulatory RNA–protein feedback loop
- Non-coding isoforms of SR genes accumulate in the nucleus under dark conditions
- Co-expression of the non-coding isoform rescues splicing factor overexpression phenotypes

## Introduction

Gene expression in eukaryotes is a complex process involving multiple layers of regulation. RNA processing, modification, and export represent crucial steps beyond transcription that fine-tune gene expression. Among these, alternative splicing (AS) is a major mechanism that generates multiple mature transcripts from a single gene, thereby greatly expanding the coding and regulatory potential of eukaryotic genomes. In plants, AS is tightly regulated by environmental conditions, including light, which is a key determinant of growth, development, and adaptation (Zhang *et al*., 2023). Our group has previously shown that light modulates AS through retrograde signals from chloroplasts, thereby impacting nuclear gene expression (Petrillo *et al*., 2014). Recently, Riegler *et al*. (2021) demonstrated that light-derived photosynthates, acting via the TOR kinase pathway, can coordinate AS changes even in non-photosynthetic tissues such as roots, revealing a systemic mechanism through which light and metabolism converge to shape splicing decisions across the whole plant.

It is commonly considered that a large proportion of genes that are subjected to AS encode proteins with regulatory roles, and the existence of AS impacts protein activity and properties. Interestingly, AS can occur through different types of events such as intron retention, exitron splicing, exon skipping, alternative 5′ or 3′ splice site selection, each of which can generate distinct transcript isoforms, different regulatory sequences, or even lead to isoforms with low or null coding potential. In plants, intron retention represents the most prevalent AS event (Marquez *et al*., 2012; Zhang *et al*., 2017, 2023), and frequently produces transcripts containing premature termination codons (PTCs). While PTC-containing transcripts are typically degraded through nonsense-mediated mRNA decay (NMD), many intron-retention isoforms including NMD features are not sensitive to this degradation pathway (Kalyna *et al*., 2012). NMD constitutes an important post-transcriptional regulatory mechanism that shapes transcript abundance. Notably, many intron-retention isoforms accumulate in the nucleus (Göhring *et al*., 2014; Petrillo, 2023), and in several cases the retained introns can be removed upon environmental or physiological cues—an “intron detention” mechanism—allowing rapid production of the fully spliced protein-coding isoforms. Although PTC-containing transcripts have traditionally been considered unproductive by-products of splicing regulation, increasing evidence indicates that non-coding isoforms generated through AS of coding genes may fulfill regulatory roles, contributing an additional layer of gene expression control (Shi *et al*., 2026).

lncRNAs are defined as transcripts longer than 200 nucleotides that lack protein-coding capacity but share several features with mRNAs, like being transcribed by RNA polymerase II and undergo 5’ cap addition, splicing, polyadenylation, and chemical modifications (Chorostecki *et al*., 2023; Saha *et al*., 2025; Zhang *et al*., 2025). Unlike mRNAs, lncRNAs typically exhibit low expression levels, with highly specific profiles of tissue, developmental stage, or environmental condition, suggesting highly specialized regulatory functions (Fonouni-Farde *et al*., 2021). Another central feature is that lncRNAs act as interaction nodes, since they can bind to proteins as well as other RNAs or DNA, thus modulating processes such as transcription, chromatin organization, and splicing. Finally, epitranscriptomic modifications, such as methylation in m6A, can influence their stability, nuclear localization, and ability to interact with proteins, reinforcing their role as dynamic regulators of gene expression (Meyer and Jaffrey, 2014; Roundtree *et al*., 2017). Interestingly, several intron-less lncRNAs were shown to interact with splicing factors or their mRNA targets to modify *in trans* specific AS patterns and the outcome of the eukaryotic transcriptome (Romero-Barrios *et al*., 2018; Rigo *et al*., 2020).

The selection of AS sites is mediated by *trans*-acting splicing factors, such as serine/arginine-rich (SR) proteins, highly conserved RNA-binding proteins. These RNA-binding proteins participate in splice site recognition and spliceosome assembly, and many of them are themselves regulated by AS and post-translational modifications such as phosphorylation (Morton *et al*., 2019). Beyond splicing, SR proteins contribute to mRNA nuclear export, stability, translation, genome maintenance, and even chromatin organization, underscoring their multifunctionality nature in the coordination of gene expression (Zheng *et al*., 2020; Wagner and Frye, 2021). In plants, the SR protein family is notably expanded compared to animals; for instance, in *Arabidopsis thaliana* this group is composed of 18 members, grouped in six sub-families with three of them being plant-specific (Barta *et al*., 2010). This diversification reflects the central role of SR proteins in modulating AS in response to developmental cues, biological timing, and biotic and abiotic stresses, thereby fine-tuning gene expression programs that support plant adaptation (Staiger and Brown, 2013).

*At-RS31* belongs to the plant-specific RS subfamily. Its protein levels increase in the light as a result of alternative splicing regulated by light–dark transitions. In response to light, *mRNA1*, the only isoform that is translated into the SR protein, increases. The other main isoforms, *mRNA2* and *mRNA3*, lack coding capacities. Notably, *mRNA2* is subjected to NMD, whereas *mRNA3*, despite sharing the same PTC, accumulates at high levels in darkness (Petrillo *et al*., 2014). Interestingly, plants overexpressing *mRNA1* (*mRNA1ox*) exhibit unfavorable phenotypes, in contrast to those overexpressing the genomic construct, which retains the native exon-intron structure and thus the capacity for AS and production of the alternative isoforms, and do not display these phenotypes, suggesting a functional role for the non-coding isoforms. We recently reported that At-RS31 protein can bind its own transcripts (Köster *et al*., 2025). In addition, the production of these non-coding isoforms in orthologs of *At-RS31* is conserved from green algae to angiosperms (Kalyna *et al*., 2006). Hence, we hypothesize that the *mRNA3* isoform, retained in the nucleus, is not a passive by-product of AS but rather acts as a regulatory RNA capable of binding and titrating splicing regulators, including its own gene product, to modulate AS in response to light/dark transitions. Interestingly, overexpression of *mRNA3* restores the wild-type phenotype to *mRNA1ox* in trans, revealing a novel paradigm in which coding genes can also give rise to regulatory lncRNAs, qualitatively widening AS transcriptome output in eukaryotes.

## Materials and Methods

### Plant material and growth conditions

The *Arabidopsis thaliana* ecotype Columbia (Col-0) was used as the wild-type in all experiments. Seeds were surface sterilized for 10 min in 30% (v/v) bleach, followed by three washes with sterile water. Seeds were stratified for three days in the dark at 4°C and then germinated on Murashige and Skoog (MS) medium buffered to pH 5.8 with 2-(N-morpholino)ethanesulfonic acid (MES) and containing 1.5% (w/v) agar. Plants were grown in a growth chamber at 21°C under long-day conditions (16-h light/8-h dark) or under continuous light at an irradiance of 70–100 μmol photons m⁻² s⁻¹. Growth conditions differing from these are specified in the text and figure legends.

### Generation of transgenic plants expressing *At-RS31* overexpression constructs

Genomic DNA and total RNA were extracted from *Arabidopsis thaliana* Col-0, and the different *At-RS31* isoforms were amplified, isolated, and purified. Isoform-specific primers used for amplification are listed in Supplementary Table S1. Each isoform was cloned into the pGreenII0029-2×35S-TL (#1068) overexpression vector using restriction enzyme–based cloning with *NcoI* and *NotI*. The resulting constructs were introduced into *Agrobacterium tumefaciens*, and Col-0 plants were transformed using the floral dip method (Clough and Bent, 1998). Transgenic seedlings were selected on MS medium supplemented with 50 μg mL⁻¹ kanamycin for 10 days, after which resistant plants were transferred to soil. For each construct, at least three independent T1 lines were obtained and subsequently advanced to homozygous T4 generations.

To generate double-overexpression lines, the *mRNA3*-p1068 construct was transformed into the stable *mRNA1ox#4* line (Petrillo *et al*., 2014). Transformants were selected under the same kanamycin/MS conditions, and at least three independent double-transgenic lines were propagated to homozygosity.

In addition, the *mRNA1* isoform was cloned into a *GreenGate* destination vector fused to GFP, following the standardized *GreenGate* cloning system (Lampropoulos *et al*., 2013).

Splicing reporter vectors were also generated using #1068 vector to express partial sequences of *At-RS31* (genomic) and *mRNA3* (partial cDNA version), with focus on the studied alternative splicing event (intron 2), and using the first intron as a control of constitutive splicing. Sequences up to the first 6 bases of exon 3 of *At-RS31* were cloned in-frame with a C-terminal hygromycin resistance cassette from pER8 using sequential assembly PCRs and restriction enzymes for cloning, generating *RS31–hyg* and *mRNA3–hyg* vectors. The sequence of *At-RS31* was mutated to eliminate all possible PTCs. Primers used for cloning and the whole strategy are documented in the Supplementary Data S1 file.

### Transient expression in *Nicotiana Benthamiana*

One-month-old *N. benthamiana* plants were grown from seed in a growth chamber at 24°C under a 16-h photoperiod using Growmix® Multipro™ substrate. *Agrobacterium tumefaciens* strains carrying the constructs of interest were cultured in liquid LB medium supplemented with kanamycin (50 µg mL⁻¹), rifampicin (100 µg mL⁻¹) and gentamicin (25 µg mL⁻¹) for 48 h at 28°C with shaking.

Bacterial cells were collected by centrifugation and resuspended to an OD₆₀₀ of 0.4 in agroinfiltration buffer (10 mM MES, 10 mM MgCl₂, and 200 μM acetosyringone). The suspension was incubated for 2 h in the dark with gentle shaking. Agroinfiltration was performed on the abaxial side of leaves using a needleless syringe, infiltrating approximately 1 mL per leaf. Plants were maintained under light conditions for 48 h prior to harvest.

### Light and drug treatments

After two weeks of growth under continuous light, plants were incubated in darkness for 48 h and then either transferred to light for 4 h or maintained in darkness for an additional 4 h.

For pharmacological treatments, seedlings grown on agar plates were submerged in 2 mL of liquid MS-MES medium supplemented with the indicated drug or dimethyl sulfoxide (DMSO) as a mock control, following the 48-h dark treatment. Vacuum infiltration was applied for 5 min to facilitate compound uptake. The drugs and final concentrations used were 2 μM or 20 μM cycloheximide (CHX; Sigma) and 100 μg mL⁻¹ actinomycin D (Act-D; Fermentek). Following treatment, seedlings were incubated under light or dark conditions for an additional 4 h.

### RNA extraction and subcellular fractionation

Total RNA was extracted using TriPure reagent (Sigma) according to the manufacturer’s instructions. Nuclear and cytoplasmic RNA fractions were isolated as described in Rodríguez *et al*. (2025).

### RT -PCR and -qPCR

For cDNA synthesis, 1 μg of RNA was reverse transcribed using the Reverse Transcription System (Sigma) with oligo(dT) primers following the manufacturer’s instructions. PCR amplification was performed using T-Plus Free DNA Polymerase (Inbio Highway) with 2 μL of 1:4 diluted cDNA. The PCR conditions were as follows: 95°C for 3 min; 30–33 cycles of 95°C for 30 s, 58–60°C for 30 s, and 72°C for 90 s; followed by a final extension at 72°C for 5 min. RT-PCR products were visualized by agarose gel electrophoresis, stained with ethidium bromide, and detected using a UV transilluminator. Band intensities were quantified by densitometry using ImageJ.

RT-qPCR analyses were performed using GreenLight Master Mix qPCR 2X (Inbio Highway) on an Eppendorf Mastercycler Realplex system.

Primer sequences are listed in Supplementary Tables S2 and S3.

### Phenotype analyses

Flowering time was assessed by counting the number of rosette leaves at bolting. Plants were grown under long-day conditions (16 h light/8 h dark) at 21°C. For each genotype, 3–6 plants per biological replicate were analyzed.

For hypocotyl length measurements, seeds were surface-sterilized, stratified for 2 days at 4°C, exposed to white light (∼100 μmol m⁻² s⁻¹) for 6 h to induce germination, and then transferred to complete darkness. Seedlings were grown vertically on half-strength MS medium containing 1.5% agar and MES for 6 days. Hypocotyl length was measured from digital images using ImageJ (20–25 seedlings per genotype per replicate).

Primary root length was measured in seedlings grown vertically on MS medium containing 1.5% agar and MES for 10 days under standard growth conditions. Root length was quantified using ImageJ (20–25 seedlings per genotype per replicate).

Root growth dynamics was analyzed using the ChronoRoot platform (Gaggion *et al*., 2021, 2026). Seeds were stratified for 2 days and grown under long-day conditions on square plates containing half-strength MS medium with 0.8% agar. Four seeds were plated per plate and eight biological replicates per genotype were analyzed.

### RNA immunoprecipitation (RIP) *in vivo* and *in vitro* assays

RIP *in vivo* assays using *RS31::RS31-GFP* lines were performed as described previously (Zhao *et al*., 2018; Rigo *et al*., 2020). Ten-day-old seedlings were harvested and cross-linked either with 1% formaldehyde for 10 min or by UV irradiation (three pulses at 0.400 J cm⁻², 254 nm UV-C, UVITEC Cambridge). RNA–protein complexes were immunoprecipitated using mouse IgG (Cell Signaling Technology) or anti-GFP antibodies (Abcam 290) at 4°C for 1 h with rotation. Dynabeads Protein A (Invitrogen) were used to capture immunocomplexes. Cross-linking was reversed and RNA was extracted using TRIzol reagent.

For *in vitro* RIP assays, total protein extracts were prepared from *RS31::RS31-GFP* seedlings grown under light or dark conditions. Nuclear protein extracts were incubated 1 hour at 4°C with *in vitro*–transcribed RNA corresponding to individual *RS31* isoforms (*mRNA1*, *mRNA2*, *mRNA3*) or with a mixture containing all isoforms. For each incubation we put 100ng of RNA of interest (individual or mixed) and 10x other RNA (GFP or tRNA) to block non-specific binding. Following the incubation, RNA–protein complexes were immunoprecipitated using GFP-TRAP antibodies (Chromotek). After extensive washing, an aliquot was taken for input (10%) and the rest was treated with proteinase K for next RNA extraction using TRIzol reagent. Enriched RNA species were subsequently analyzed by RT-PCR or RT-qPCR using isoform-specific primers. Primers used for *in vitro* transcribed isoforms are listed in Supplementary Table S4.

### Statistical analyses

Statistical analyses were performed using GraphPad Prism Software version 8.0.2 for Windows (GraphPad Software, San Diego, CA, USA) or InfoStat Software version 2017 (Grupo InfoStat, FCA, Universidad Nacional de Córdoba, Argentina). Data were analyzed by analysis of variance followed by Tukey’s or Fisher’s least significant difference (LSD) post hoc test. Different letters or * indicate statistically significant differences (p < 0.05).

### Plant material for RNA sequencing

Four *Arabidopsis thaliana* lines were analyzed: *mRNA1ox #4* (Petrillo *et al*., 2014), *mRNA3ox #15*, the double overexpression line *mRNA(1+3)ox #4A2*, and wild-type (WT, Col-0). Plants were grown under the same conditions used for ChronoRoot experiments. Three biological replicates per genotype were analyzed, each consisting of two 10-day-old seedlings collected from independent plates.

### AtRTD2 PTC detection

PTC detection is based on STOP codons presence upstream of a splicing junction (>50 nt rule). We implemented a streamlined three-step pipeline in Python to perform the analyses. Using the original AtRTD2_19April2016 fasta (.fa) and gtf (.gtf) files (available at https://ics.hutton.ac.uk/atRTD/) we generated an enriched FASTA with CDSs and Junctions with the “add_junction.py” script. Since AtRTD2 has many isoforms from coding genes that lack any associated CDS, and also some annotated isoforms with shifted START codons (compared to others in the same gene), we analyzed the transcriptome gene-by-gene to re-assign proper and common START codons to all the isoforms, using the script “reannotate_CDS.py”. The enriched and corrected FASTA was then used to detect the PTCs with the script “detect_ptcs.py”.

### RNA extraction and library preparation

Total RNA was extracted using the protocol above. RNA samples were treated with TURBO™ DNase (Life Technologies) according to the manufacturer’s instructions to remove residual genomic DNA and subsequently purified using the RNA Clean & Concentrator™ kit (Zymo Research). RNA concentration and quality were assessed prior to library preparation. For RNA sequencing, 600 ng of total RNA per sample were used to generate poly(A)-selected libraries using the NEBNext® Ultra™ II Directional RNA Library Prep Kit (New England Biolabs), following the manufacturer’s instructions, resulting in libraries with an average fragment size of 200–500 bp. Library quality and size distribution were assessed using an Agilent Bioanalyzer with the High Sensitivity DNA Kit (Agilent). A second cleanup step was performed using SPRIselect beads (Beckman Coulter) to remove residual adapter contamination. Indexed libraries were pooled and sequenced by Novogene.

### RNAseq read alignment

Raw reads were quality-assessed with FastQC (www.bioinformatics.babraham.ac.uk/projects/fastqc/). Then, clean reads were mapped to the TAIR10 genome with STAR aligner (version 2.5.2b) using a one-pass mapping with the following options: --readFilesCommand zcat, --outSAMtype BAM SortedByCoordinate, --quantMode TranscriptomeSAM GeneCounts, --alignSJoverhangMin 8, --alignSJDBoverhangMin 1, --alignIntronMin 20, --alignIntronMa× 10000, --alignMatesGapMa× 10000, --outFilterMismatchNoverLmax 0.04, --outFilterMultimapNma× 10, --outFilterScoreMinOverLread 0.66, --outFilterMatchNminOverLread 0.66. The resulting BAM files were indexed with SAMtools for proper visualization of reads and splice junctions in Integrative Genome Viewer (IGV). Sashimi plots were used to visualize splicing junction counts in IGV. To this end, the threshold for SetJunctionCoverageMin was set to 5 for every track.

### 3D RNA-seq app analyses

This section of the Method details was adapted from the output “Results” of the 3D RNA-seq package (Guo *et al*., 2019). Transcript per million (TPM) normalization was estimated using Salmon with option --validateMappings. Salmon output (.sf files) was used as input for 3D RNAseq App using ATRTD2_QUASI reference transcriptome since it covers most of the known alternatively spliced isoforms in Arabidopsis thaliana (Patro *et al*., 2017; Zhang *et al*., 2017).

Read counts were generated using tximport R package version 1.10.0 and lengthScaledTPM method (Soneson *et al*., 2016) with inputs of transcript quantifications from Salmon. Low-expressed transcripts and genes were filtered based on analyzing the data mean-variance trend. A gene/transcript was considered expressed if it had at least two counts per million (CPM ≥ 2) in at least three samples The TMM method was used to normalize the gene and transcript read counts to log2(CPM) (Bullard *et al*., 2010).

The Limma R package with its voom function was used for differential gene expression, splicing and transcript expression comparison (Law *et al*., 2014; Ritchie *et al*., 2015). Differential gene expression (DE) and transcript usage (DTU), log2 (Fold Change) of gene/transcript abundance were calculated based on contrast groups and significance of expression changes were determined using t test. P values of multiple testing were adjusted with Benjamini–Hochberg (BH) to correct false discovery rate (FDR) (Benjamini and Hochberg, 1995). A gene/transcript was significantly DE in a contrast group if it had an adjusted p-value < 0.05 and |log2(FC) ≥ 1|. At the alternative splicing level, DTU transcripts were determined by comparing the log2(FC) of a transcript to the weighted average of log2(FC) (weights were based on their standard deviation) of all remaining transcripts in the same gene. A transcript was determined as significant DTU if it had an adjusted p-value < 0.05 and ΔPercent Spliced (ΔPS) ≥ 0.1. For differentially alternatively spliced (DAS) genes, each individual transcript log2(FC) was compared to gene level log2(FC), which was calculated as the weighted average of log2(FC) of all transcripts of the gene. Then p values of individual transcript comparison were summarized to a single gene level p value with F-test. A gene was significantly DAS in a contrast group if it had an adjusted p value < 0.01 and any of its transcript had a ΔPS ratio ≥ 0.1.

### Venn Diagrams

Tables of significantly affected genes obtained from the 3D RNA-seq tool analyses were the input for Venny 2.1 (An interactive tool for comparing lists with Venn’s diagrams, developed by Oliveros JC, https://bioinfogp.cnb.csic.es/tools/venny/index.html).

### Heat Map plots

Gene expression across all genotypes was assessed with Heat Map plots. To avoid noisy transcript expression, genes with TPM ≥ 10 in all replicates were kept using a custom python script. Also, to construct the heat map, genes with a fold change > 2 (relative to Col-0) were used. The expression matrix was scaled to a Z-score matrix and the k-means function of R was used for gene clustering with option set.seed(123) for reproducibility.

Differential splicing heat map was constructed using the ΔPS of each genotype (relative to Col-0) using a custom python script with seaborn and matplotlib libraries. To ensure splicing changes, if a gene ΔPS had a p-adj. value > 0.05, it was automatically assigned a value of zero.

Splicing index heat map of SR genes to evaluate phenotype reversion of mRNA(1+3)ox line was constructed by calculating splicing index (longest isoform TPMs divided by the sum of TPMs of all isoforms). Such splicing indexes were plotted using a custom Python script with seaborn and matplotlib libraries.

## Results

### 1. Alternative splicing generates abundant PTC-containing isoforms that accumulate in the nucleus in *Arabidopsis thaliana*

Alternative splicing contributes to transcriptome complexity in plants, yet its contribution to protein diversity is often limited, as intron retention is the most frequent event. In addition, transcripts with NMD features often accumulate to high levels and, in some cases, they escape degradation by accumulating in the nucleus (Kalyna *et al*., 2006; Göhring *et al*., 2014; Petrillo *et al*., 2014; Fuchs *et al*., 2021). To assess the prevalence of transcripts that possess NMD triggering features in *Arabidopsis thaliana*, we interrogated its transcriptome (AtRTD2) with a custom-made pipeline (https://github.com/petrylab/PTC-Analysis) for AS isoforms containing PTCs. Strikingly, more than half (52.24%) of the genes with alternative isoforms harbor at least one PTC-containing isoform (Figure 1A). Moreover, as the number of isoforms per gene increases, so does the likelihood of generating a PTC-containing variant (Figure 1B), resulting in a reduced proportion of non-PTC, most likely, protein-coding isoforms (Figures 1C–D). Thus, *A. thaliana* AS frequently generates nonproductive transcripts with features that could trigger NMD. This phenomenon is particularly pronounced in SR protein–coding genes, which display multiple PTC-containing isoforms that accumulate substantially in RNA-seq datasets (Supplementary Data S2) and RT-PCR assays (Palusa *et al*., 2007; Palusa and Reddy, 2010; Petrillo *et al*., 2014; Hartmann *et al*., 2018). Remarkably, certain SR protein-genes such as *At-RS31* and *At-SR30* show extreme accumulation of nonproductive isoforms, reaching over 60% and 80% of total transcripts, respectively, under dark-grown seedling conditions (Supplementary Data S2).

**Figure 1:**
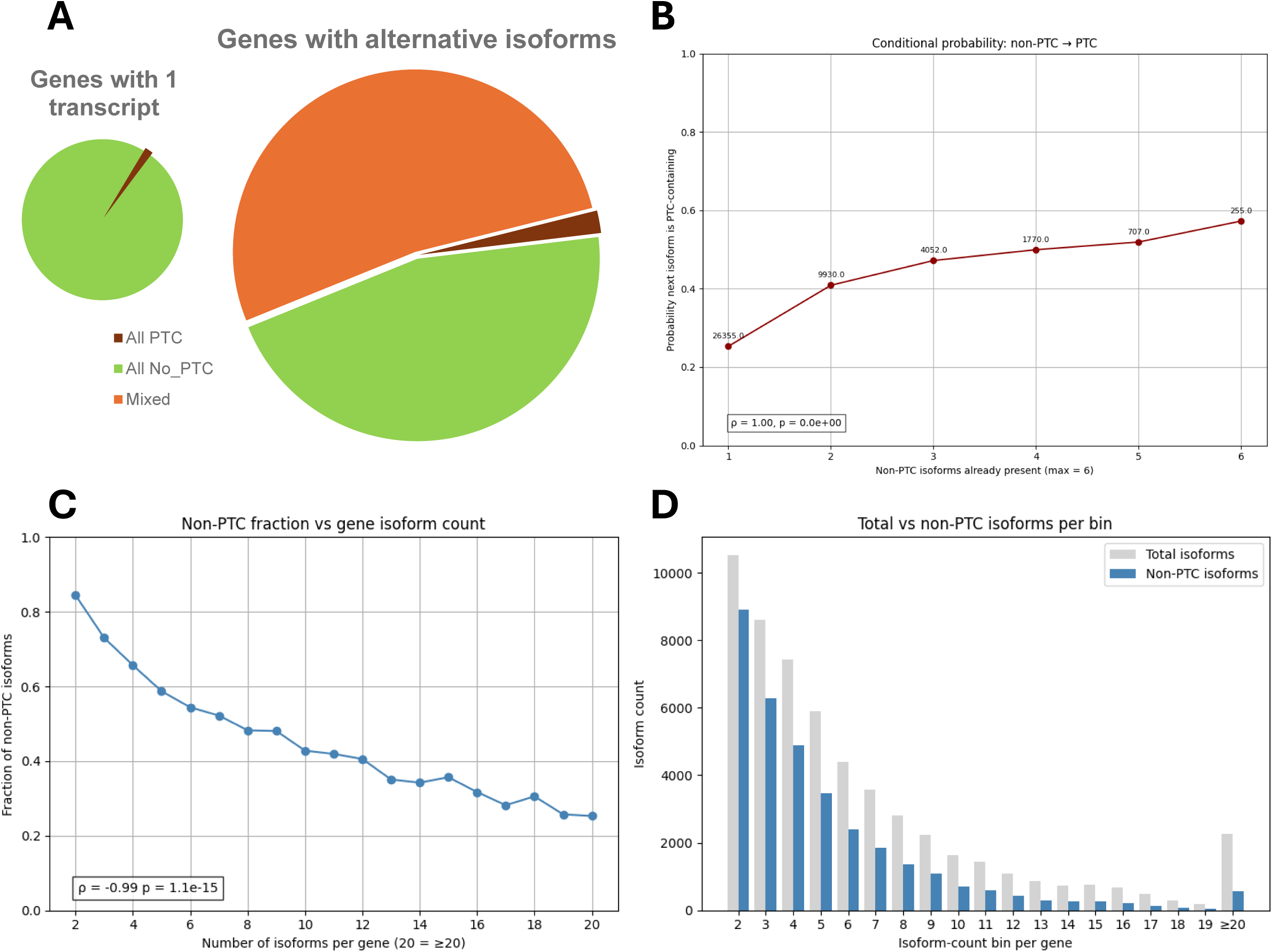
Increasing isoform diversity correlates with diminishing coding capacity in *Arabidopsis thaliana*. **(A)** Isoforms annotated in the AtRTD2 transcriptome were evaluated for the presence of premature termination codons (PTCs). To standardize coding potential, all isoforms were re-annotated to initiate translation from the same, most likely AUG start codon. PTCs were defined as stop codons located >50 nucleotides upstream of an exon–exon junction. The pie chart displays the distribution of genes with a single annotated isoform (left) and genes with alternative isoforms (right), subdivided into those where all isoforms are predicted to contain a PTC (dark red), all are non-PTC containing (green), or a mixture of PTC and non-PTC containing isoforms (orange). A substantial fraction of multi-isoform genes (52.24%) produces at least one PTC-containing transcript. **(B)** The probability that an isoform contains a premature termination codon (PTC) was calculated conditional on the number of coding isoforms already present in a gene. The graph shows that the likelihood of generating a PTC-containing isoform increases as isoform diversity expands. A positive and significant Spearman correlation supports this trend, indicating that genes with more coding isoforms are more prone to produce PTC-containing transcripts. Numbers on top of the points are the genes considered. **(C)** Scatter plot showing the fraction of productive isoforms relative to the number of isoforms per gene. Genes with higher numbers of isoforms tend to have a lower proportion of productive transcripts. Spearman correlation supports this trend. **(D)** Bar chart comparing the total number of isoforms (gray) and the number of productive isoforms (blue) across bins of isoform counts per gene. Bins are coincident with isoform number from 2 to 19, but from 20 onwards, all bins are collapsed. Productive isoforms consistently represent a smaller subset of the total, with the disparity becoming more pronounced in genes with higher isoform counts.

We hypothesized that these PTC-containing isoforms can persist by accumulating in the nucleus, as previously shown for some members of the SR protein gene family (Göhring *et al*., 2014; Hartmann *et al*., 2018; Fuchs *et al*., 2021). Using our recently developed *SuB3* fractionation protocol (Rodríguez *et al*., 2025) combined with RT–PCR AS analysis we observed that most SR protein-coding genes produce at least two isoforms, with one isoform enriched in the nuclear fraction (Figure 2A). To gain deeper insight into these regulatory mechanisms, we analyzed the well characterized AS events of *At-SR30* and *At-RS31*, aiming to assess the coding capacities and subcellular distribution of their different isoforms. Furthermore, both splicing index measurements and RT–qPCR quantification of specific isoforms revealed a progressive accumulation (relative and absolute) of the non-coding isoforms of these genes as the duration of darkness incubation increased (Figure 2B and 2C).

**Figure 2.**
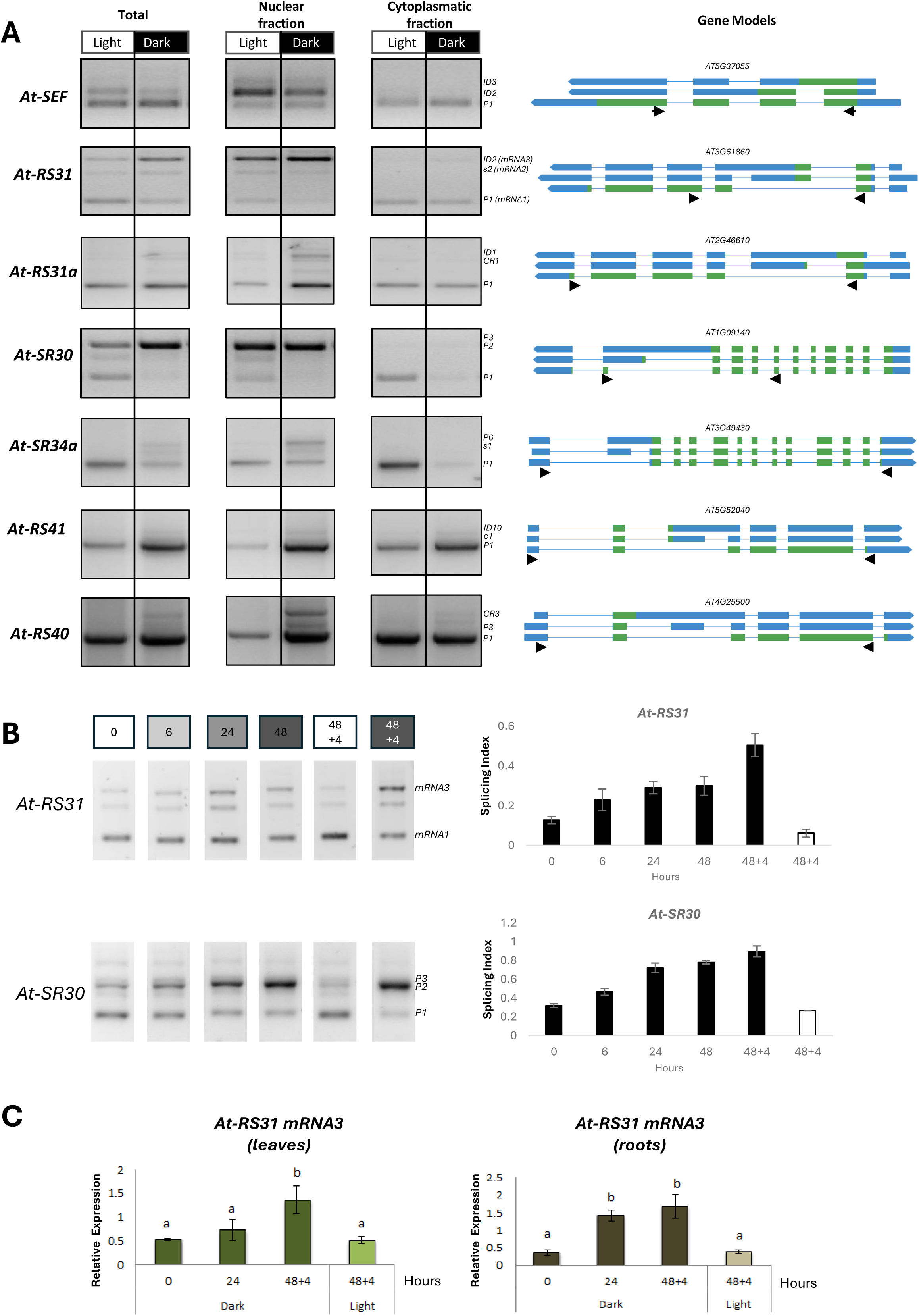
Light-dependent nuclear accumulation of non-coding SR transcript isoforms. **(A)** Alternative splicing patterns from Total, Nuclear-and Cytoplasmic-enriched fractions obtained using the *SuB3* protocol (Rodríguez *et al*., 2025) from *Arabidopsis thaliana* seedlings exposed to light (Light) or darkness (Dark), were analyzed by RT–PCR using specific primers for each SR gene (see Supplementary Table S2). SEF was used as a fractionation control. Gene model and all alternative isoforms are illustrated from Boxify (boxify.boku.ac.at/). **(B)** Splice variants of *At-RS31* and *At-SR30* were analyzed by RT-PCR during the dark-light transition, and each PCR product was quantified using the Splicing *Index*. **(C)** RT–qPCR quantification of non-coding transcript abundance for At-RS31 relative to PP2A as a housekeeping gene in leaves and in roots. Bars represent mean ± SD (n≥3). Significant differences are shown with different letters (p<0.05 by ANOVA & Tukey’s post-test).

Together, these findings suggest that light/dark transitions modulate the AS of SR protein–coding genes primarily by shifting transcript output toward non-coding variants that accumulate in the nucleus, rather than substantially diversifying protein isoforms, and this nuclear enrichment may limit their degradation by NMD.

### 2. *At-RS31* isoform balance controls light-responsive splicing and development

Given the stability and high relative levels of accumulation of the non-coding isoforms generated by AS of SR coding genes, we next focused on *At-RS31*, a highly regulated AS event in light/dark transitions, to explore the potential functions of those non-coding nuclear transcripts. *At-RS31* undergoes AS, producing three major isoforms with distinct coding capacities (Figure 3A). *mRNA1* corresponds to the coding transcript; *mRNA2*, which includes an alternative exon that generates a PTC, is NMD-sensitive; and *mRNA3*, which includes the same alternative exon plus a small intronic sequence (sub-intron), is stable and nuclearly retained (Fig. 2A and Petrillo *et al*., 2014). Furthermore, plants overexpressing the coding isoform *(mRNA1ox*) display growth defects in normal growth conditions, while plants overexpressing a genomic construct (*31GenOX*) show an enhanced growth phenotype, compared with the wild-type (Col-0) (Petrillo *et al*., 2014). The splicing pattern of *At-RS31* is regulated by light via chloroplast-derived retrograde signals (Petrillo *et al*., 2014) and several mRNA targets of At-RS31 protein have been identified using iCLIP (Köster *et al*., 2025). In darkness, the *mRNA3* steadily increases, while light exposure rapidly reverses the splicing pattern favoring the *mRNA1* isoform (Fig. 2C–D). To assess the functional impact of these isoforms, we analyzed *At-RS31* AS pattern in transgenic lines overexpressing *mRNA1* (*mRNA1ox #4 and #14*) and compared them with wild-type, *31GenOX*, and *rs31-1* mutant plants. Overexpression of *mRNA1* markedly altered the splicing pattern of the endogenous copy of *At-RS31* itself and that of several downstream targets including other SR genes such as *At-SR30*, *At-RS40*, and *At-SCL30a* (Fig. 3B and 3E). Hence, At-RS31 splicing factor is regulating the AS outcomes of its own gene and also influencing the isoforms relative abundance of several members of the SR protein family.

**Figure 3.**
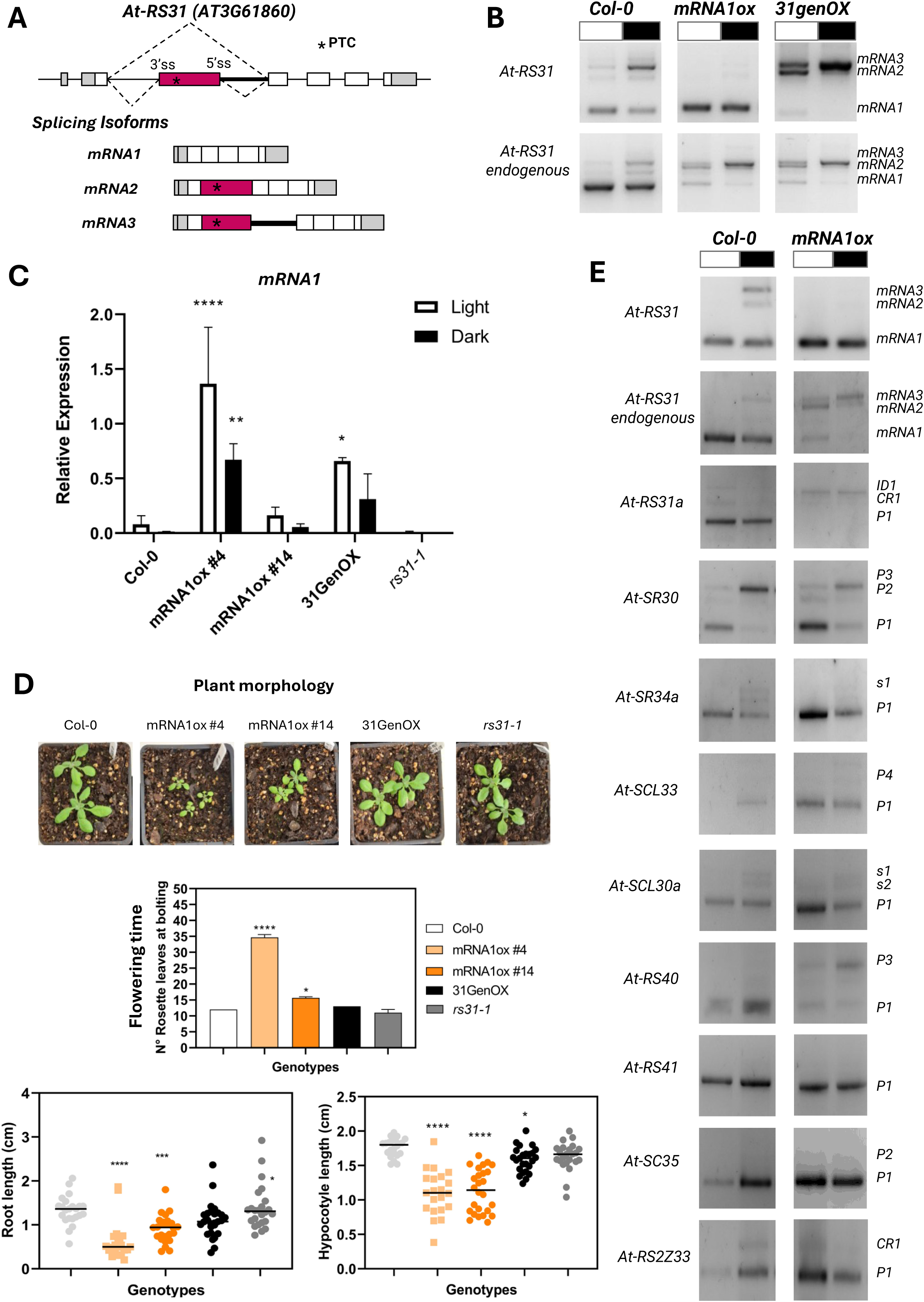
Functional analysis of *At-RS31* isoforms. **(A)** Schematic representation of the *At-RS31* gene structure and its three main splicing isoforms. **(B)** Alternative splicing patterns of *At-RS31* from seedlings of different lines (Col-0, *mRNA1ox* and *31GenOx*) grown under light (white box) or dark (black box) conditions were analyzed by RT–PCR using two specific sets of primers: one that detects both the transgene and the endogenous gene (*At-RS31*) and another one that detects only the endogenous gene (*At-RS31 endogenous*), see the Supplementary Table S2. **(C)** RT–qPCR quantification of *At*-*RS31* in all the lines (Col-0, *mRNA1ox #4* and *#14, 31GenOx,* and *rs31-1* mutant). *mRNA1* expression levels were normalized to *PP2A*. Bars represent mean ± SD (n=3). * indicates significant differences (p<0.05) relative to Col-0 by ANOVA & Tukey’s post-test. **(D)** Phenotypic analysis of the different At-RS31 de-regulated plants where changes in plant morphology were observed. Number of rosette leaves at bolting, Primary root length(cm) of 10-day-old seedlings and Hypocotyl length (cm) of 6-day-old seedlings. Bars represent mean ± SD (n≥5, n≥20, n≥20). Significant differences are shown with * p<0.05 by ANOVA & Tukey post-test. (E) RT–PCR validation of SR genes affected by *mRNA1ox* and At-RS31 targets identified by iCLIP. Primers are listed in the Supplementary Table S2.

Phenotypically, *mRNA1ox* plants exhibited pronounced developmental defects, including reduced rosette size, delayed flowering (also reported in Saile *et al*., 2025), and altered root and hypocotyl elongation under dark conditions (Figure 3D). RT–qPCR analysis confirmed the overexpression of *At-RS31* in both *mRNA1ox* lines and in the *At-RS31* genomic overexpression line (*31GenOX*) and revealed differential expression across the different transgenic lines (Figure 3C). Interestingly, despite the fact that the *mRNA1ox* lines exhibit different expression levels, they consistently reproduce the developmental phenotypes (Figure 3D and also observed Saile *et al.,* 2025). In contrast, the *At-RS31* genomic overexpression line, which also displays high levels of the coding isoform *mRNA1*, shows a wild-type-like phenotype. The difference with the *mRNA1ox* overexpression line is that the *31GenOX* also exhibits high levels of expression of the other isoforms, i.e. *mRNA3* and *mRNA2* (Figure 3B). This suggests that the presence of these isoforms, though devoid of coding capacity, is functionally relevant to modulate At-RS31 splicing factor function.

### 3. Post-transcriptional conversion of At-RS31 mRNA3 into the NMD-sensitive mRNA2

In light of these results, we next sought to characterize the non-coding isoforms (*mRNA2* and *mRNA3*) in greater detail to explore their potential roles. Under normal conditions, *mRNA2* is degraded through the nonsense-mediated mRNA decay (NMD) pathway as evidenced by the stabilization of this transcript upon treatment with cycloheximide (CHX), an inhibitor of translation and, concomitantly, NMD (Figure 4A and Supplementary Figure S1). Interestingly, when overexpressing the *mRNA1* or the *At-RS31* genomic construct, there is a substantial increase of the *mRNA2* isoform (Figure 3B). RT-PCR analysis using endogenous primers revealed that *mRNA2* accumulates to some extent in the nucleus of *mRNA1*-overexpressing lines, which may explain its higher stability (or accumulation) in this transgenic line (Figure 4B). In addition, relative levels of *mRNA2* are higher in light than in darkness (Figure 4B and 4C), hinting at a light-mediated regulation of this isoform. To properly assess the accumulation kinetics of the *mRNA2*, we performed a time-course analysis on NMD-defective *upf3-1* mutants, revealing an increase of *mRNA2* accumulation after 90 minutes of light exposure (Figure 4D). This behavior coincides with the detected reversion of the splicing pattern of *At-RS31* in wild-type plants (Petrillo *et al*., 2014 and this work), suggesting that the generation of *mRNA2* is temporarily linked to the splicing shift.

**Figure 4.**
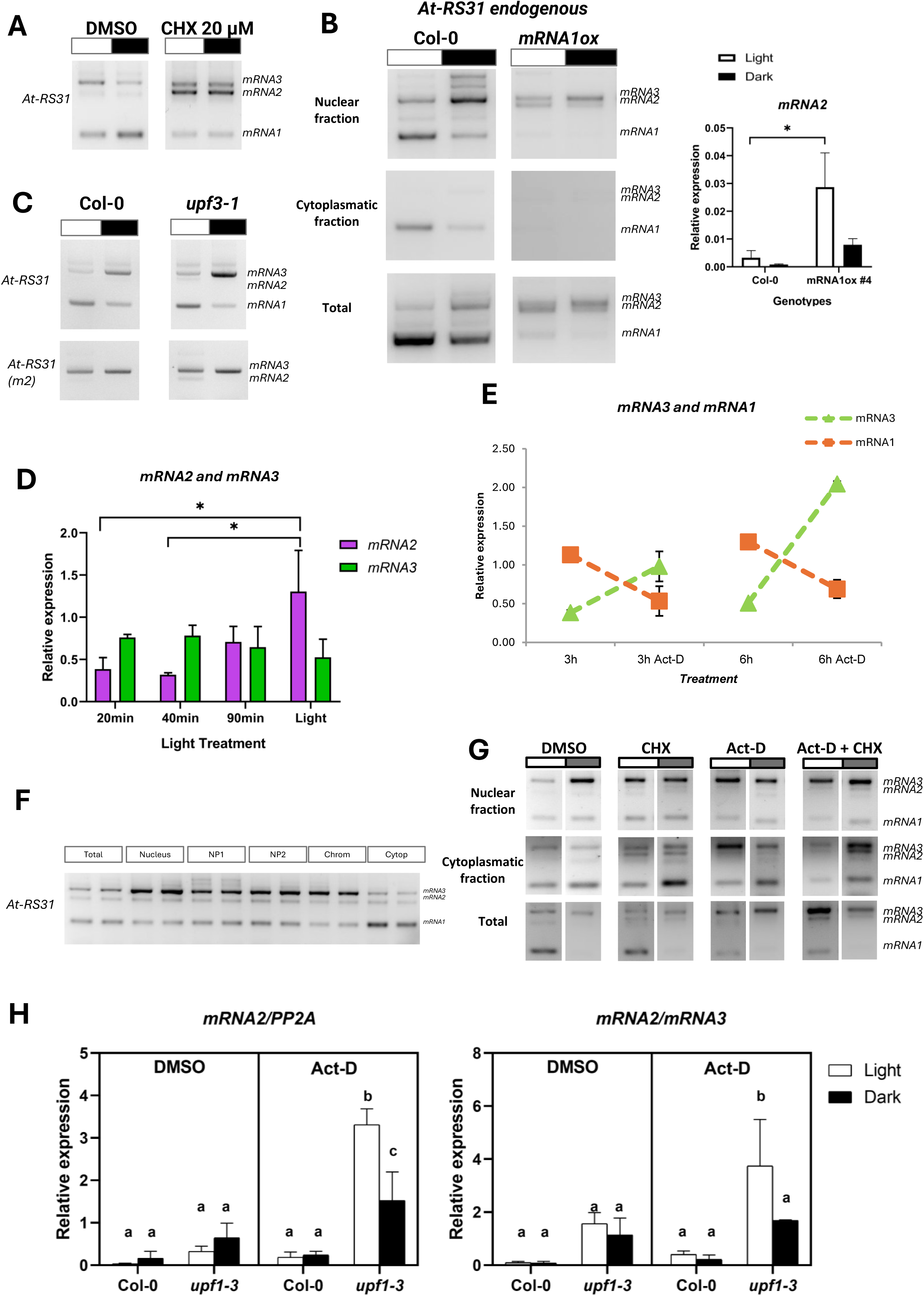
Characterization of *At-RS31* non-coding isoforms and their regulation by NMD and light. **(A)** Alternative splicing patterns of *At-RS31* from Col-0 seedling treated with CHX (20 µM) under light or dark conditions were analyzed by RT–PCR using specific primers. **(B)** Left, alternative splicing patterns from Total RNA, and Nuclear- and Cytoplasmic-enriched fractions obtained using the *SuB3* protocol (Rodríguez *et al*., 2025) from *Arabidopsis thaliana* seedlings of both genotypes (Col-0 and *mRNA1ox*) exposed to light or darkness. RT–PCR analysis was carried out using specific primers that detects only the endogenous gene (*At-RS31* endogenous). Right, RT–qPCR quantification of *At*-*RS31* in the same lines. *mRNA2* expression levels were normalized to *PP2A.* Bars represent mean ± SD (n=3). * indicates significant differences (p<0.05) relative to Col-0 by ANOVA & Bonferroni’s post-test. **(C)** Alternative splicing patterns of Col-0 and NMD-deficient *upf3-1* mutant line in light and dark. **(D)** RT–qPCR quantification of *At-RS31* in NMD-deficient mutant (*upf3-1*) grown in different light time exposure (20, 40 and 90 minutes) or always in light. *mRNA2* and *mRNA3* expression levels were normalized to *PP2A.* Bars represent mean ± SD (n=3). * indicates significant differences (p<0.05) relative to Col-0 by ANOVA & Tukeys’s post-test. **(E)** RT–qPCR quantification of *At-RS31* in Col-0 seedlings treated with actinomycin D after 3 or 6 hours. *mRNA1* and *mRNA3* expression levels were normalized to *UBI.* **(F)** Alternative splicing patterns from Total RNA, and Nuclear, Nucleoplasm 1 (NP1), Nucleoplasm 2 (NP2), Chromatin (Chrom) and Cytoplasmic (Cytop)-enriched fractions obtained using the *SuB3* protocol from Col-0 seedlings were analyzed by RT–PCR using primers for *At-RS31* splicing. **(G)** Alternative splicing patterns from Total RNA, and Nuclear- and Cytoplasmic-enriched fractions obtained using the *SuB3* protocol from *Arabidopsis thaliana* seedlings of Col-0 seedlings treated with cycloheximide (CHX) and/or actinomycin D (Act-D), grown under light or dark conditions were analyzed by RT–PCR using specific primers (see Supplementary Table S2). **(H)** RT–qPCR quantification of *At-RS31* in Col-0 or *upf3-1* treated with Act-D, grown in light or dark condition. *mRNA2* expression levels were normalized to *PP2A* or to *mRNA3.* Bars represent mean ± SD (n=3). Different letters indicate significant differences (p<0.05) by ANOVA & Sidak’s post-test.

In contrast, *mRNA3* accumulates in the nucleus and preferentially under dark conditions (Petrillo *et al*., 2014). We confirmed that *mRNA3* is stable in light and dark, as its levels do not preferentially decrease upon 3-6 hours of actinomycin D (Act-D) transcriptional inhibition assays (Figure 4E). Interestingly, though it has similar features to *mRNA2*, the nuclear accumulation of *mRNA3* allows it to evade NMD. Since this could be interpreted as the *mRNA3* is a non-fully processed transcript that remains attached to the polymerase or it is not released from the transcription site, we evaluated if *mRNA3* accumulates in the nucleus as a chromatin associated RNA. However, we found this isoform in both chromatin-bound and nucleoplasmic fractions. Hence, the *mRNA3* exists as a freely diffusing nuclear RNA (Figure 4F and Figure 4G).

Together, these results indicate that *mRNA2* functions as a canonical NMD-sensitive isoform, whereas *mRNA3* represents a stable, nuclear-retained transcript. Notably, the temporal dynamics of *mRNA2* accumulation in NMD-deficient lines, combined with the splicing pattern shift observed in wild-type seedlings, the nuclear localization of *mRNA3*, and its sequence features, suggest that *mRNA2* may arise post-transcriptionally from *mRNA3* splicing. Coherently, RT-PCR analysis of *At-RS31* intron 2 splicing identifies *mRNA3* as the predominant detectable intermediate, supporting a stepwise pathway in which removal of the upstream intron 2 segment (sub-intron) precedes the excision of the downstream *mRNA3* sub-intron (Supplementary Figure S2). To dissect whether *mRNA2* arises from post-transcriptional processing rather than *de novo* transcription, we treated *Arabidopsis* seedlings with Act-D and CHX, separately or in combination (Figure 4G). While Act-D treatment led to accumulation of *mRNA3* and CHX induced *mRNA2* accumulation, the combined treatment resulted primarily in *mRNA2* enrichment. This supports the hypothesis that *mRNA2* is derived from *mRNA3* through an *At-RS31*-dependent post-transcriptional splicing event. Finally, to further confirm the involvement of the NMD pathway, we analyzed *upf3-1* mutant plants treated with Act-D and observed a significant light-dependent increase in *mRNA2* levels, particularly under illumination (Figure 4H). Remarkably, this post-transcriptional splicing mechanism offers a means to eliminate *mRNA3* from the nucleus upon light exposure, yet it also raises the question of why this transcript would accumulate in the first place.

### 4. The At-RS31 protein selectively binds *mRNA3 in vitro* and *in vivo*

The observations of nuclear accumulation of *mRNA3* in darkness and its depletion under light conditions raised the possibility of a veiled function for this transcript. To elucidate its possible role, we generated *mRNA3* overexpression lines (*mRNA3ox*) in the wild-type background. The *mRNA3*-overexpressing plants did not display any obvious phenotypic differences compared to wild-type plants under any of the physiological conditions tested, nor did they show detectable changes in the AS patterns of known *At-RS31* target genes (Supplementary Figure S3). Interestingly, plants overexpressing the genomic *At-RS31* construct (*31GenOX*), which display high expression levels of both *mRNA1* and *mRNA3*, do not exhibit the phenotypic defects observed in *mRNA1ox* lines, particularly those regarding primary root length and flowering time (Figure 3 and Supplementary Figure S3).

These findings imply that the *mRNA3* plays a functional role specifically in the context of *At-RS31* overexpression, potentially modulating the effects associated with elevated At-RS31 protein levels. In this scenario, the *mRNA3* could be titrating At-RS31 protein and diminishing its availability to affect its targets. Supporting this idea, previous iCLIP data revealed that the At-RS31 protein binds to its own transcripts (Supplementary Figure S4, Köster *et al*., 2025), although the specific isoform or precursor molecule involved could not be resolved. This observation led us to test whether *mRNA3* may exert a regulatory role through a direct interaction with the At-RS31 protein. Then, we performed RNA immunoprecipitation (RIP) assays using *RS31::RS31-GFP* transgenic lines to evaluate whether *mRNA3* is among the RNAs bound by At-RS31 *in vivo*. Remarkably, both under light and dark conditions, *mRNA3* was the most enriched isoform in the immunoprecipitated fraction (Figure 5A). However, given that *in vivo* RIP assays could potentially co-purify nascent pre-mRNAs still associated with chromatin, we sought to confirm these findings using an *in vitro* approach. For this purpose, we purified *RS31::RS31-GFP* protein from *A. thaliana* plants and incubated it with *in vitro*–transcribed isoforms of *At-RS31*, either individually or in combination (Figure 5B-C). Consistent with the *in vivo* results, the *in vitro* RIP assays showed that *mRNA3* was preferentially bound by At-RS31 protein, displaying the highest enrichment.

**Figure 5.**
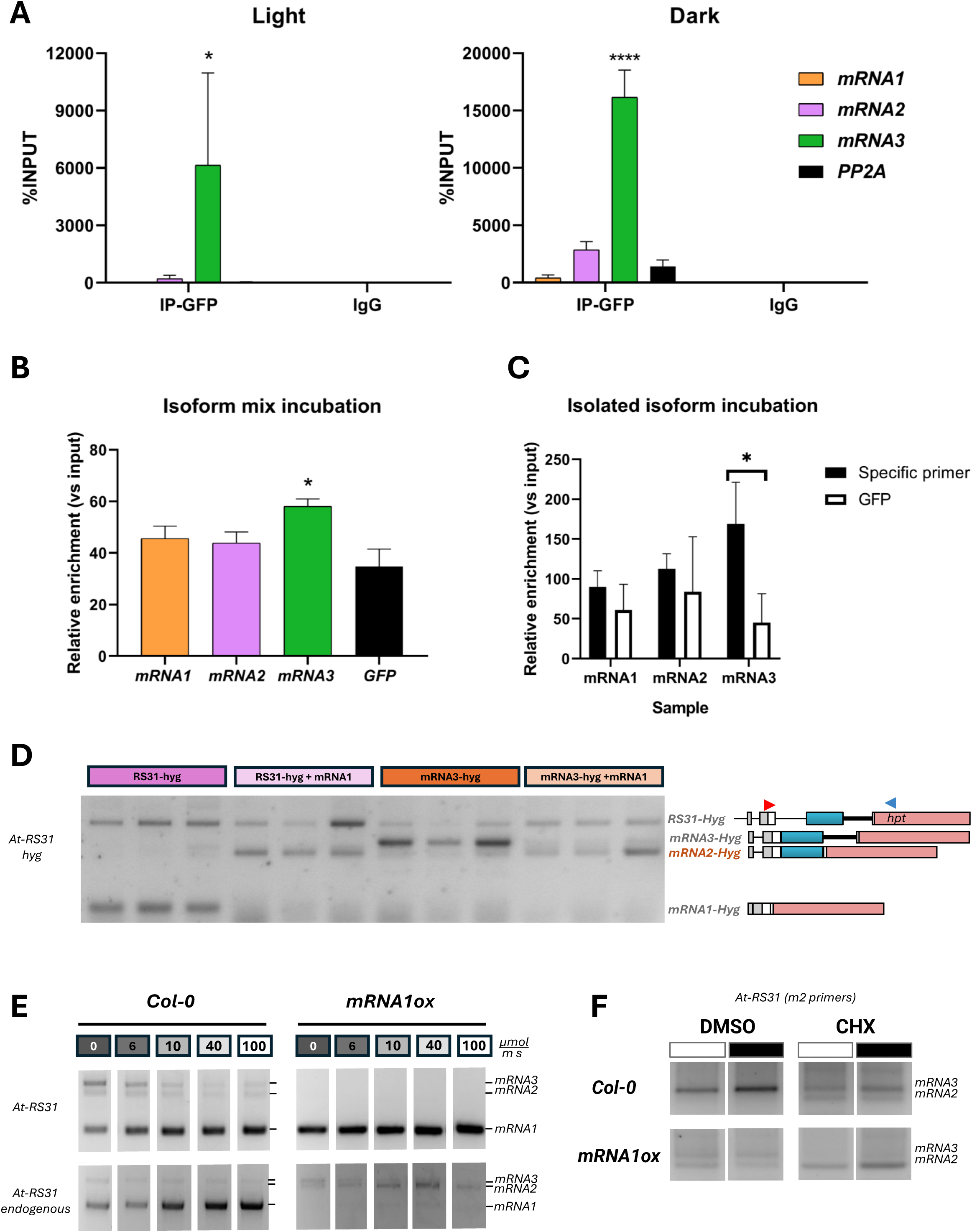
At-RS31 protein preferentially binds to its non-coding isoform *mRNA3* and enhances its post-transcriptional splicing into *mRNA2*. **(A)** *In vivo* RNA immunoprecipitation (RIP) assays were performed using *RS31::RS31-GFP* transgenic *Arabidopsis thaliana* lines grown under light or dark conditions. RT–qPCR analysis of *At-RS31* isoform expression levels from immunoprecipitated (IP) or IgG relative to Input fractions. Bars represent mean ± SD (n=2). * indicates p<0.05 by ANOVA & Bonferroni test. **(B)** I*n vitro* RIP assays were performed using *RS31::RS31-GFP* protein purified from *Arabidopsis thaliana* plants (in light or in darkness) and incubated with *in vitro*–transcribed *At-RS31* isoforms (*mRNA1, mRNA2,* and *mRNA3*), either in combination **(B)** or separately **(C).** RT–qPCR analysis of *At-RS31* isoform expression levels from immunoprecipitated (IP) or IgG relative to Input fractions. Expression was normalized to *GFP* in C. Bars represent mean ± SD (n=2). * indicates p<0.05 by ANOVA & Bonferroni test (square-root transformed data was used). **(D)** Alternative splicing patterns of *At-RS31* from *Nicotiana benthamiana* leaves transient expression assays with *RS31-hyg* or *mRNA3-hyg* reporter constructs co-infiltrated with or without *mRNA1*-GFP vector were analyzed by RT–PCR using specific primers (RS31spl Fw and HYG Rv). **(E)** Analysis of alternative splicing by RT–PCR of *At-RS31* endogenous transcripts in Col-0 and *mRNA1ox* seedlings exposed to different light intensities (0, 6, 10, 40 and 100 μmol m⁻² s⁻¹) using two specific sets of primers: one that detects both the transgene and the endogenous gene (*At*-*RS31*) and the other one that detects only the endogenous gene (*At-RS31* endogenous). **(F)** Alternative splicing patterns from Col-0 and *mRNA1ox* seedlings treated with CHX 20 µM under light or dark conditions were analyzed by RT–PCR using specific primers to detect *mRNA2* or *mRNA3* (see Supplementary Table S2).

Our findings indicate that At-RS31 selectively recognizes and binds the *mRNA3* isoform. This prompted us to evaluate if the interaction could have a processing consequence for this isoform. We designed chimeric reporter constructs: one containing the first exons and introns (1 and 2) of *At-RS31* (*RS31-hyg*), and another one as a partial cDNA construct in which intron 1 is conserved but intron 2 is replaced with the sequence of the *mRNA3* isoform (*mRNA3-hyg*). These constructs were transiently expressed in *Nicotiana benthamiana* leaves. Importantly, neither construct can produce At-RS31 protein, as they lack the coding sequence from exon 3 onwards. Furthermore, they are not subject to NMD, since all potential PTCs were eliminated. To evaluate the effect of At-RS31 protein on their splicing behavior, the constructs were infiltrated either alone or co-infiltrated with an *mRNA1* construct encoding At-RS31 protein (Figure 5D). In the absence of At-RS31 protein, the *RS31hyg* construct predominantly produces the equivalent of the coding *mRNA1* isoform. By contrast, co-expression of *mRNA1* to give the At-RS31 protein, shifted the splicing pattern of *RS31hyg* towards the equivalent of the *mRNA2* isoform. A similar outcome was observed with the *mRNA3* reporter, which is spliced into the *mRNA2*-like variant upon At-RS31 co-infiltration.

These findings indicate that At-RS31 binding promotes the splicing of *mRNA3* into *mRNA2*, thereby functionally linking the two non-coding isoforms in a dynamic autoregulatory loop. To determine whether this mechanism operates in its natural cellular environment, we analyzed *mRNA1ox A. thaliana* plants exposed to different light intensities. RT–PCR analysis of the endogenous *At-RS31* gene revealed that *mRNA1* overexpression enhances *mRNA2* accumulation. Furthermore, increasing light intensity correlated with elevated *mRNA2/mRNA3* ratios (Figure 5E). These effects are more dramatic when abolishing NMD, using CHX treatment *mRNA2* levels outcompete *mRNA3* in a semi-quantitative RT-PCR (Figure 5F). Together, these results support a model in which At-RS31 protein regulates the turnover of *mRNA3* via splicing to generate *mRNA2*, which is subsequently degraded by NMD. We hypothesize this mechanism finely tunes At-RS31 protein levels in response to light/dark transitions and, in turn, *mRNA3* can act as a decoy RNA contributing to temporarily titrating At-RS31 protein. Such a mechanism may explain why the *31GenOX* lines do not show the phenotypic effects associated with excess accumulation of At-RS31.

### 5. The *mRNA3* counterbalances high At-RS31 protein levels *in trans*

Having established that the At-RS31 protein promotes the turnover of *mRNA3* by splicing it into the NMD-sensitive isoform *mRNA2*, we next asked whether *mRNA3* alone can regulate At-RS31 function or, more specifically, its downstream gene-expression effects. To address this question, we generated and transformed the original *mRNA1ox* transgenic line to overexpress the *mRNA3* as well, generating double overexpression lines (*mRNA(1+3)ox*). To verify correct transgenes expression, we performed RT–qPCR analysis, confirming the presence of both *mRNA1* and *mRNA3* transcripts in the double overexpression lines (Figure 6A). Although *mRNA1* levels were slightly lower than those of the strongest *mRNA1ox* line (#4), they were comparable to those of line #14, which nevertheless displayed clear *mRNA1* overexpression phenotypes. Consistent with these observations, the splicing pattern of *At-RS31* in *mRNA(1+3)ox* plants resembled that of the *31GenOX* line (Figure 6B).

**Figure 6.**
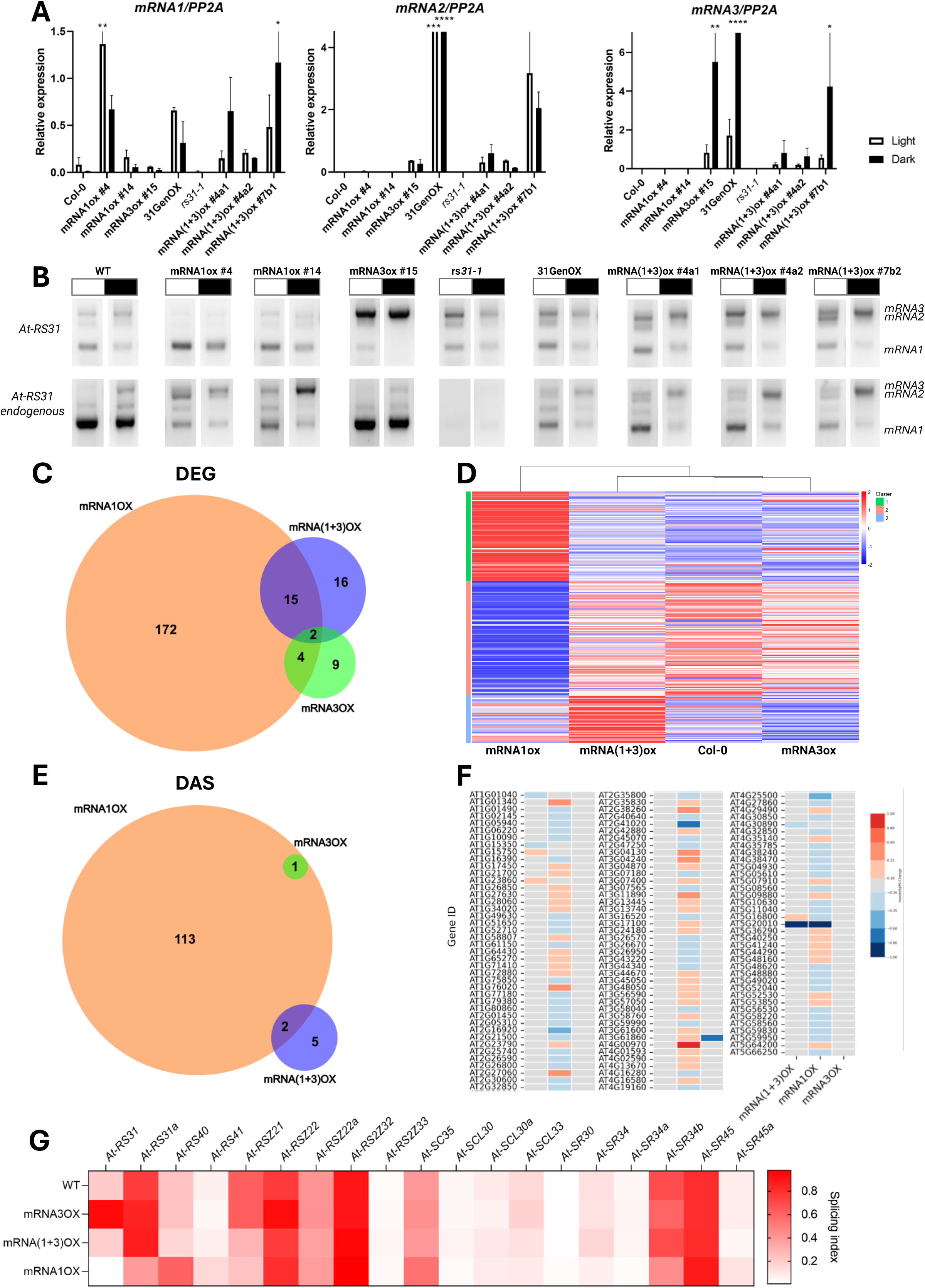
Characterization of molecular phenotype of *mRNA(1+3)ox* line. **(A)** RT–qPCR analysis of *At-RS31* isoform expression levels in Col-0, *mRNA1ox* lines, *mRNA3ox* lines, *mRNA(1+3)ox* lines and *rs31-1* grown under light or dark conditions. Expression of each isoform was normalized to *PP2A*. Bars represent mean ± SD (n=3). * indicates p<0.05 by ANOVA & Fisher’spost-test. **(B)** Alternative splicing patterns of *At-RS31* from seedling of different transgenic lines (Col-0, *mRNA1ox #4 and #14, mRNA3ox #15, rs31-1 and mRNA(1+3) #4a1, #4a2 and #7b2)* grown under light or dark conditions were analyzed by RT–PCR using two specific sets of primers: one that detects both the transgene and the endogenous gene (*At-RS31*) and the other one that detects only the endogenous gene (*At-RS31* endogenous) see Supplementary Table S2 for further details. **(C)** Venn Diagram showing the overlap among differentially expressed genes (DEG) in *mRNA1ox, mRNA3ox* and *mRNA(1+3)ox* lines obtained by RNAseq analysis. **(D)** Differentially expressed genes (DEG) in *mRNA1ox, mRNA3ox* and *mRNA(1+3)ox* lines obtained by RNAseq analysis were visualized by a heatmap of K-means clustering of z-scores of the means of normalized reads (TPMs). Hierarchical clustering of the genotypes is shown as a dendrogram. **(E)** Venn Diagram showing the overlap among differentially spliced genes (DAS) in *mRNA1ox, mRNA3ox* and *mRNA(1+3)ox* lines obtained by RNAseq analysis. **(F)** Differentially spliced genes (DAS) in *mRNA1ox, mRNA3ox* and *mRNA(1+3)ox* lines obtained by RNAseq analysis were visualized by a heatmap showing the ΔPS. Red: ΔPS > 0.1 and blue ΔPS < −0.1. **(G)** Splicing index (SI) of *SR* genes in *mRNA1ox, mRNA3ox* and *mRNA(1+3)ox* lines obtained by RNAseq analysis were visualized by a heatmap.

Expectedly, the double overexpression lines show high levels of *mRNA2* accumulation, which evidence that both *mRNA1* and *mRNA3* are present in high levels while reinforcing the notion that At-RS31 protein enhances *mRNA3* splicing into *mRNA2*. Hence, we further investigated the transcriptome profiles of the single-isoform overexpression lines, *mRNA1ox* and *mRNA3ox*, as well as that of the double overexpression line, *mRNA(1+3)ox*, to assess whether the presence of *mRNA3* is able to buffer or modulate the widespread transcriptional changes triggered by At-RS31 (i.e.: *mRNA1* overexpression) (Köster *et al*., 2025). Interestingly, although *mRNA1* overexpression alone alters the expression of numerous genes, the vast majority of these changes are absent when *mRNA1* is co-overexpressed with *mRNA3* (Figures 6C–D and Supplementary Table S5). Similarly, the genes whose AS patterns are affected in *mRNA1ox* plants are largely unchanged in the *mRNA(1+3)ox* lines (Figures 6E–F and Supplementary Table S6), underscoring a broad buffering role of *mRNA3* at both the transcriptional and co-/post- transcriptional levels. Remarkably, while the AS pattern of the endogenous *At-RS31* in the *mRNA(1+3)ox* is different than that of the wild-type plant and resembles the regulation observed in the *31GenOX* line, the double overexpression line mostly mirrors the splicing indexes observed in wild-type plants for other regulated SR protein coding genes (Figure 6G).

Considering that *mRNA1* overexpression causes deleterious phenotypes while *At-RS31* genomic construct overexpression does not, and that *mRNA3* interacts directly with the At-RS31 protein and molecularly restores gene expression patterns, we next asked whether the overexpression of this non-coding isoform in a context of high protein levels would rescue the developmental defects associated with *mRNA1* overexpression. Interestingly, several of the deleterious phenotypes observed in *mRNA1ox* plants were partially or completely reverted in the double overexpression lines. In particular, flowering time and hypocotyl length in darkness returned to wild-type levels (Figure 7A). Using the automated root phenotyping system *ChronoRoot* (Gaggion *et al*., 2021, 2026), we further observed that *mRNA(1+3)ox* plants displayed an intermediate root growth phenotype compared with wild-type and *mRNA1ox* plants, supporting a partial restoration of normal root growth dynamics by the co-expression in *trans* of the *mRNA3* isoform (Figure 7B).

**Figure 7:**
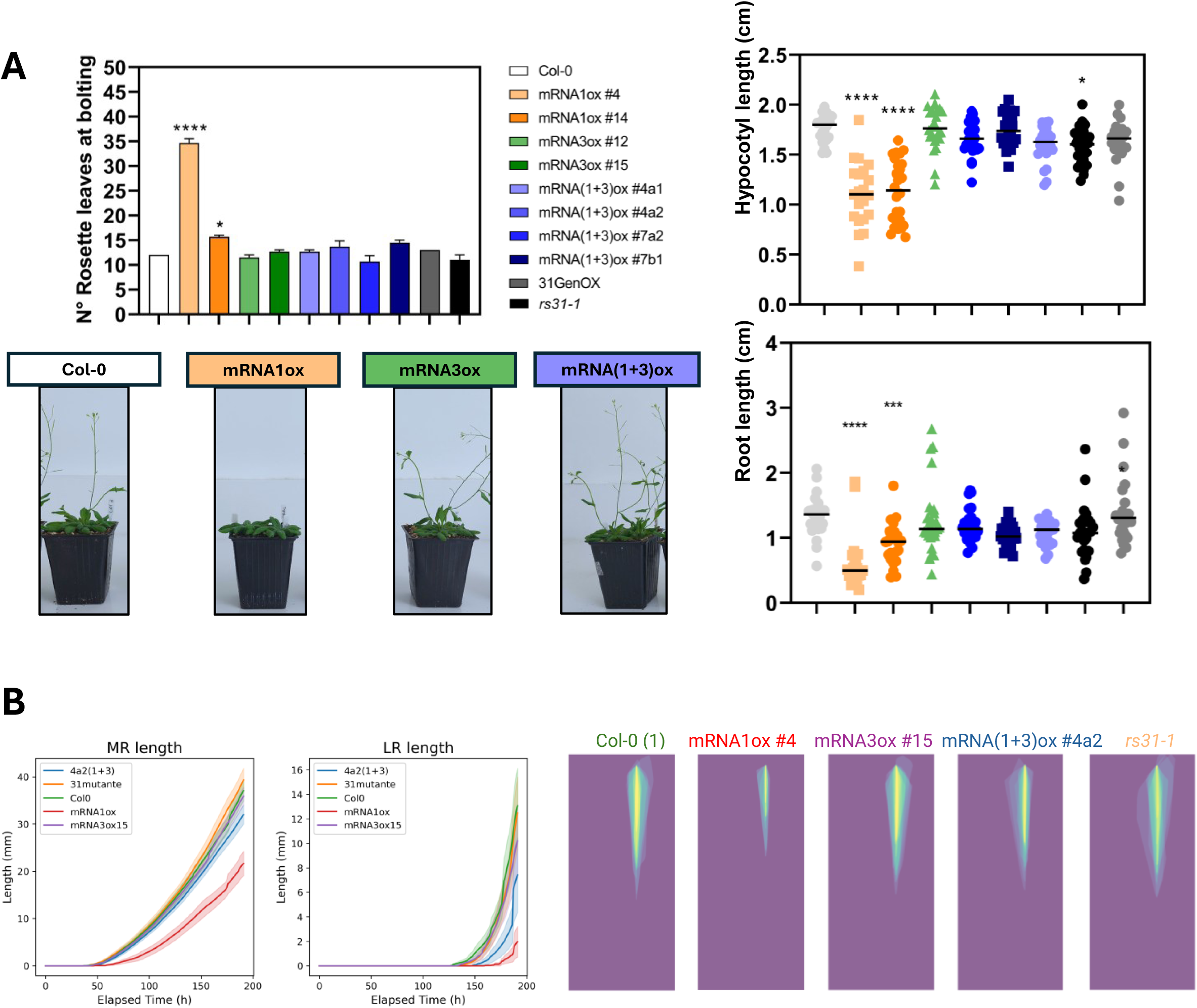
Overexpression of the non-coding isoform *mRNA3* in the high-level *mRNA1-*overexpression background partially rescues the deleterious phenotypes of *At-RS31* overexpression. **(A)** Phenotypic characterization of double overexpression lines (*mRNA1+3ox*) compared with wild-type (Col-0), *mRNA1ox*, *mRNA3ox*, *31GenOX* and *rs31-1* plants: Flowering time (Number of rosette leaves at bolting (left panel) and the corresponding plant images (down panel), primary root length(cm) of 10-d-old seedlings and hypocotyl length (cm) of 6-day-old seedlings (right panel). Each bar represents the mean ± SD (n≥3 flowering, n≥20 roots, n≥20 hypocotyls). Statistically significant differences relative to Col-0 between means are indicated by * (p<0.05, one-way ANOVA followed by Dunnett’s post-test). **(B)** Comparison of root growth parameters in seedlings grown for 8 days on MS medium under long day conditions. Average primary root length (mm) and root architecture were evaluated for each genotype using ChronoRoot (Gaggion *et al*., 2026).

Our findings build on established knowledge on AS regulation and support a model in which the coding gene *At-RS31* produces both a protein-coding isoform (*mRNA1*) and a nuclear non-coding isoform that feeds back on At-RS31 activity. In this framework, *mRNA3* acts as a decoy RNA that engages At-RS31 protein subsequently promoting its own post-transcriptional splicing into the NMD-sensitive isoform *mRNA2*, helping maintain isoform balance and splicing factor homeostasis. This dual output modulates At-RS31-dependent transcriptome changes and, consequently, plant development and light-dark transitions. More broadly, lncRNAs derived from protein-coding loci may enrich the complex regulatory mechanisms controlling the eukaryotic transcriptome in response to environmental cues.

## Discussion

AS is traditionally seen as a source of transcriptomic and ergo, proteomic diversity. In plants, however, this view is somewhat short-sighted, as intron retention is the main outcome and most resulting isoforms contain PTCs that would route them to NMD. Interestingly, a vast proportion of these transcripts can escape NMD by staying in the nuclear compartment and accumulate to relatively high levels. This raises the question whether these transcripts are simply by-products, or whether they play active regulatory roles. Darwin’s principles question why a plant would generate and accumulate a large number of seemingly non-functional transcripts only to degrade them later. Why invest energy in producing these isoforms in the first place? This paradox, showing the real face of plants as the most energetically ostentatious organisms on Earth, may suggest that these transcripts may not be useless, but instead fulfill active regulatory roles underlying adaptation and evolution.

Our data supports this latter view. Specifically, the *mRNA3* isoform that accumulates in the nucleus under dark conditions remains stable and interacts directly with At-RS31 protein, displaying features consistent with a regulatory lncRNA (Kopp and Mendell, 2018; Mattick *et al*., 2023). Alternatively, *mRNA3* could be a passive byproduct of *At-RS31* AS regulation. In this scenario its accumulation would be anecdotal. Yet the fact that At-RS31 binds this transcriptional isoform complicates such a dismissal. Its nuclear presence, even if passive, could sequester At-RS31 or subtly alter its distribution, thereby influencing regulatory outcomes. Building on this isoform-specific case, our results reveal a broader feedback mechanism that connects splicing regulation to environmental cues.

Our present results extend previous work showing that light regulates AS via chloroplast retrograde signals and TOR kinase pathways (Riegler *et al*., 2021). By uncovering a feedback mechanism in which At-RS31 protein levels are modulated by its own non-coding isoforms, we provide a mechanistic link between environmental cues (light/dark transitions), splicing regulation, and plant physiology. Together with our recent report showing At-RS31 protein can bind its own transcripts and those of other SR coding genes (Köster *et al*., 2025), our results position At-RS31 as a central node in the SR protein networks. The interplay between coding and non-coding isoforms thus contributes to the robustness and adaptability of splicing regulation under fluctuating environmental conditions. Furthermore, we demonstrate that At-RS31 protein binds preferentially to its own non-coding isoform and promotes post-transcriptional processing of *mRNA3* into *mRNA2*. This autoregulatory mechanism establishes a feedback loop that fine-tunes At-RS31 expression through the interplay of RNA–protein interactions and NMD sensitivity. Our results therefore demonstrate that SR proteins can modulate their own impact on the plant transcriptome through AS-dependent non-coding intermediates.

The light-dependent shift in *At-RS31* isoform composition provides a mechanism to buffer the availability of functional SR protein for splicing regulation, directly linking AS to environmental cues and plant adaptation. Similar phenomena have been reported in animal cells exposed to UV irradiation (Williamson *et al*., 2017), where non-coding transcripts are generated by alternative polyadenylation combined with intron retention. In wheat, the *TAVRN1 ALTERNATIVE SPLICING* (*VAS*) lncRNA is derived from the locus encoding the TaVRN1 transcription factor, as a product of AS. *VAS* physically associates with the bZIP transcription TaRF2b, inducing *TaVRN1* transcription during vernalization and accelerating flowering. Thus, *VAS* is an example of a lncRNA derived from a protein-coding locus affecting the transcriptional activity of its parental coding gene (Xu *et al*., 2021). In *A. thaliana*, it was proposed that a circular RNA originating by back-splicing from the sixth exon of the *SEPALLATA3* (*SEP3*) locus can hybridize with its own genomic sequence to generate a DNA–RNA hybrid structure. This R-loop facilitates AS of the emerging transcript, favoring production of the naturally occurring exon 6–skipped isoform, *SEP3.3*. The accumulation of this variant was linked to alterations in floral organ identity (Conn *et al*., 2017). In both flies and humans, the splicing regulator MUSCLEBLIND has been shown to bind with high affinity and specificity to a circRNA generated from its own gene, known as *circMbl* (Ashwal-Fluss *et al*., 2014). This circular RNA has been proposed to act as a competitor of the linear splicing process of its parental transcript, modulating the isoform population of the *MUSCLEBLIND* transcripts. Here, our findings highlight a transcript produced through AS which is bound by the protein encoded from the same parental gene, underscoring the role of this mechanism as a powerful amplifier of gene expression regulation. The capacity of protein-coding genes to give rise to regulatory lncRNAs through AS introduces a paradigm shift in our understanding of gene expression.

It suggests that the functional output of a gene cannot be fully captured by its protein product alone but should also account for the regulatory roles of its non-coding isoforms. This duality expands the toolkit of AS outcomes, providing plants-and likely other eukaryotes-with versatile strategies to fine-tune gene expression in response to environmental and physiological signals. Future studies should explore whether similar autoregulatory loops exist in other SR genes and whether this mechanism is conserved across plant species or even in other eukaryotes.

Beyond *At-RS31*, we also identified other transcripts with potential regulatory functions in the SR protein coding genes family. A particularly intriguing example is *At-SR30*. This gene produces multiple isoforms, with *SR30.1* encoding the protein (it is *SR30.P1* here). Hartmann *et al*. (2018) showed that *SR30.2* (*SR30.P2* here) accumulates predominantly in the nucleus and to higher levels in darkness (as we show in Figure 2). Notably, the authors also showed that *SR30.2* can undergo further splicing to generate several NMD-sensitive isoforms by using different alternative 5′ splice sites (see JBrowse and Supplementary Data S1). This suggests that precise splicing decisions are not critical in this case, what matters is their efficient export to ensure their degradation (Hartmann *et al*., 2018). Albeit these similarities, SR30.2/P2 possible regulatory roles still need to be evaluated. In this sense, and also from Wachter lab, there is intriguing evidence linked to a conserved RNA structural element, known as DEAD, that is present in the DEAD-box RNA helicase (DRH) gene family from land plants. The authors show that in *A. thaliana* this element regulates the use of an alternative splice site as part of a negative feedback loop that also relies on the helicase binding to this region (Burgardt *et al*., 2026). Remarkably, this RNA motif is present in the middle of the longest intron (the 4th) of *At-DRH1* (AT3G01540), and the alternative splicing outcomes closely resemble those of *At-RS31.* In the future, overexpressing the alternative splicing (*alt3* and *CE*) isoforms of *At-DRH1* will also help determine if the mechanism we propose is conserved in different gene families throughout the *A. thaliana* genome.

Interestingly, more than half of the genes from *A. thaliana -*according to the AtRTD2 transcriptome-that enjoy AS have PTC-containing isoforms. This observation hints at massive regulatory consequences. Yet a key consideration must be addressed: many canonical intron retention events, when stable and retained in the nucleus, may actually represent detained introns, i.e. those not immediately removed during pre-mRNA splicing. This does not preclude them from being regulatory and active, but it places them in a different category than the one we describe here. In these cases, resumption of splicing post-transcriptionally would generate the coding isoform, meaning that the retained transcripts function as reservoirs of productive mRNAs rather than as independent regulatory entities. Furthermore, nuclear RNA concentration has been proposed to directly regulate transcription in human cells (Berry *et al*., 2022). Perturbations that elevate nuclear mRNA levels trigger a compensatory downregulation of transcriptional activity, establishing a negative feedback loop that maintains RNA concentration homeostasis. This mechanism involves reduced abundance of RNA polymerase II and altered states of transcription-associated proteins, thereby coordinating RNA synthesis with nuclear mRNA degradation and export. Such findings highlight a broader principle: nuclear-retained transcripts are not merely passive intermediates but can actively influence gene expression dynamics through feedback on transcription. In light of our results, it is tempting to speculate that plant nuclear isoforms, including *mRNA3*, may contribute to analogous homeostatic regulation, either by modulating the availability of splicing factors or by shaping transcriptional output in response to environmental cues. This possibility situates our work within a growing recognition that AS-derived non-coding isoforms can act as regulatory agents, expanding the functional repertoire of the transcriptome beyond protein coding capacity.

Importantly, light signals were also demonstrated to control RNA polymerase II activity in *A. thaliana.* In this framework, Micaela Godoy Herz and colleagues (Godoy Herz *et al*., 2019) have shown that AS regulation is coupled to RNA polymerase II elongation in response to light/dark transitions. Here we demonstrated that the *mRNA3* could be further, post-transcriptionally, spliced to produce the NMD-sensitive isoform *mRNA2,* producing the AS shift upon light exposure. Our present work is adding an extra layer of complexity to the current model of *AtRS31* AS regulation, since both mechanisms, co-and post-transcriptional, are likely acting on *At-RS31* transcripts. On the one hand, light/dark conditions may regulate the choice of 3′ splice sites, determining the production of *mRNA1* or *mRNA3* in a cotranscriptional manner. On the other hand, the efficient elimination of stable nuclear *mRNA3* transcripts requires a post-transcriptional splicing step to generate the NMD-sensitive *mRNA2*. Together, these layers of regulation highlight how transcriptional and post-transcriptional processes converge to fine-tune isoform composition in response to environmental cues.

Even though it lies beyond the scope of the present work, we cannot exclude the possibility that the interaction between At-RS31 protein and the *mRNA3* isoform regulates the activity or status of the splicing factor at an additional level, such as phosphorylation. Our model is consistent with the view that At-RS31 phosphorylation, in addition to its abundance, can be modulated by light. Upon light exposure, a change in At-RS31 phospho-status could trigger the recruitment of spliceosome components to the *mRNA3*-bound transcripts, thereby activating the splicing of its remaining sub-intron. Considering that in metazoans it has been shown that the interaction of proteins with lncRNAs triggers protein post-translational modifications of splicing factors (Romero-Barrios *et al*., 2018), further research will help to decipher whether the interaction between At-RS31 and the *mRNA3* isoform regulates its activity or phosphorylation. This represents an exciting possibility that warrants future investigation, as it would integrate transcriptional, post-transcriptional, and post-translational regulation into a unified framework for light-dependent control of alternative splicing.

Light not only shapes transcript composition through AS but also exerts a profound influence on cytoplasmic translation. It was demonstrated that during early photomorphogenesis, thousands of *A. thaliana* transcripts undergo selective translational regulation, with ribosome occupancy and density markedly increasing after light exposure (Liu *et al*., 2012). This genome-wide enhancement of translation efficiency was independent of steady-state mRNA abundance and favored transcripts with specific molecular signatures, such as shorter coding sequences and defined 5′ UTR motifs. Within this framework, it is reasonable to propose that the *mRNA1* isoform of *At-RS31* would be more efficiently translated under light conditions, where its levels are also rising (Petrillo *et al*., 2014), thereby elevating At-RS31 protein levels. The combination of isoform-specific splicing regulation and light-enhanced translation thus provides a coherent mechanism by which environmental cues dynamically tune At-RS31 abundance and, in turn, splicing regulation across the SR protein network.

Thinking in absolutes (though it works for *Siths* and for politicians running the world these days) is not appropriate in biology. Labels as “coding” and “non-coding”, as many other labels, are far from fixed; they are constantly revisited in daily biological discourse and in light of new evidence. We have shown that a coding gene generates both a protein-coding isoform and non-coding RNAs, which together orchestrate a regulatory feedback mechanism. According to our proposed model (Figure 8), the non-coding *mRNA3* accumulates in darkness and it is able to titrate At-RS31 protein during the initial phase of light exposure, reducing the effective availability of At-RS31 as a splicing regulator for its other targets. As translation resumes in light, At-RS31 protein levels rise, promoting the post-transcriptional splicing of *mRNA3* into *mRNA2,* effectively switching off the titration mechanism, and closing the autoregulatory loop. This regulatory feedback loop provides a mechanism to fine-tune At-RS31 activity and maintain splicing homeostasis properly adjusting it to light/dark transitions (Figure 8). Furthermore, these results provide a framework for exploring how AS-derived lncRNAs contribute to gene regulatory networks and environmental responsiveness.

**Figure 8:**
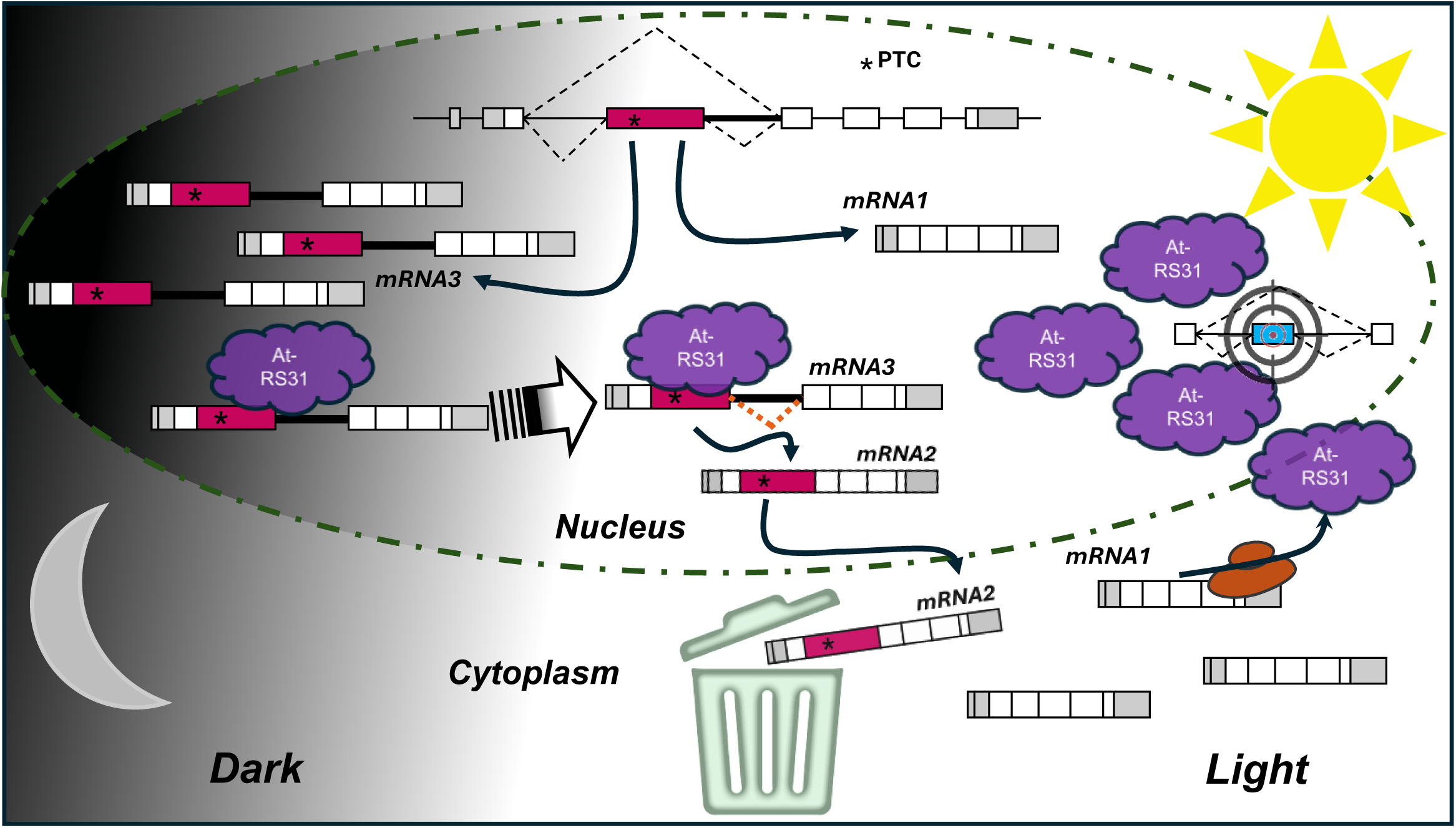
Proposed model of *At*-*RS31* alternative splicing regulation. In darkness (left panel), the non-coding isoform *mRNA3* accumulates in the nucleus and interacts with the At-RS31 protein, which is also less abundant under these conditions. This interaction may limit At-RS31 availability for binding to its other RNA targets, thereby maintaining low splicing activity. Upon illumination (right panel), light-induced signals promote the expression of the coding *mRNA1* isoform, increasing At-RS31 protein abundance. At-RS31 then binds to nuclear *mRNA3*, promoting its post-transcriptional splicing in light (as indicated by the striped arrow) into *mRNA2*, which is subsequently degraded through the NMD pathway. The *mRNA3* nuclear isoform binds to At-RS31 and buffers the excess of At-RS31. This feedback mechanism allows the system to dynamically adjust At-RS31 protein levels and activity according to light conditions, ensuring proper regulation of alternative splicing and downstream gene expression.

## Acknowledgments

The authors would like to thank all the wonderful people at IFIBYNE, Apolo, Max Perutz Labs, BOKU, and IPS2 for the fantastic atmospheres for learning, working, and progress—not only the scientific staff, but also the administrative teams and the broader communities that make our work possible and easygoing. The EP lab was supported by the ANPCyT (Agencia Nacional de Promoción Científica y Tecnológica, Argentina, PICTs: 2017-1343, 2019-01690 and 2020-02865) until funding was suspended, and it is now solely supported by ICGEB CRP/ARG22-03. The FA lab is supported by the AXA Research Fund (AXA Chair) and the RIBOLEG project (Argentina). MC and FA are supported by the IRP LOCOYM funded by the CNRS. FSR and FEA are PhD fellows, and RST, FA, and EP are career researchers from CONICET.

## Declaration of generative AI and AI-assisted technologies in the manuscript preparation process

During the preparation of this work the authors used Claude, ChatGPT, Copilot and Gemini in order to correct informatic scripts and text, to produce a clearer version for most readers. After using these tools and services, the authors fully reviewed and substantially edited the content, as needed, and take full responsibility for the content of the published article.

---

**Suppl. Figure S1.**
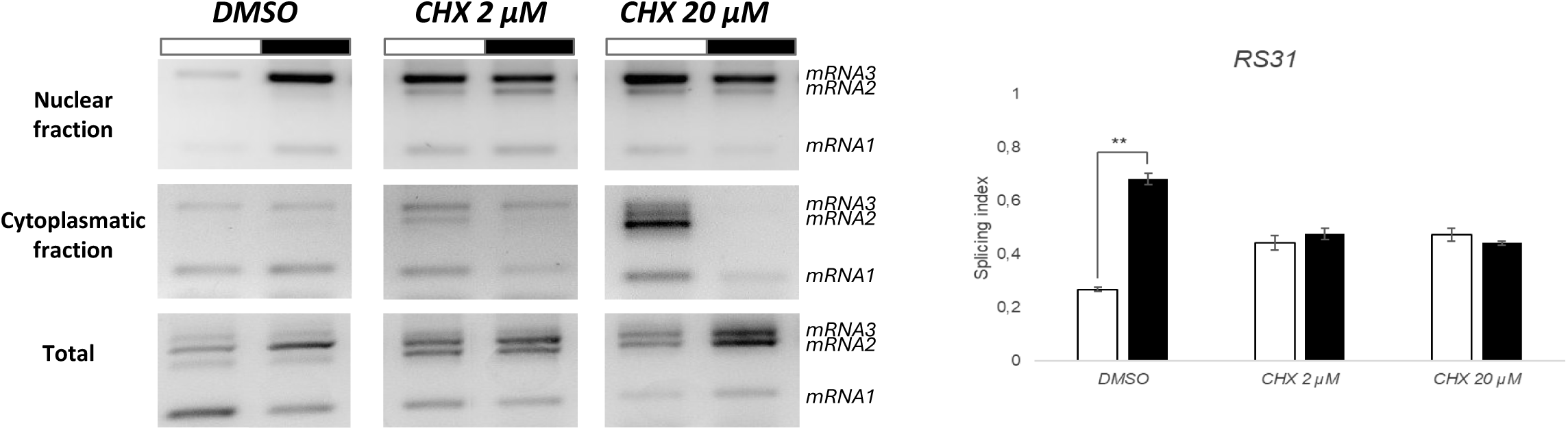

**Suppl. Figure S2.**
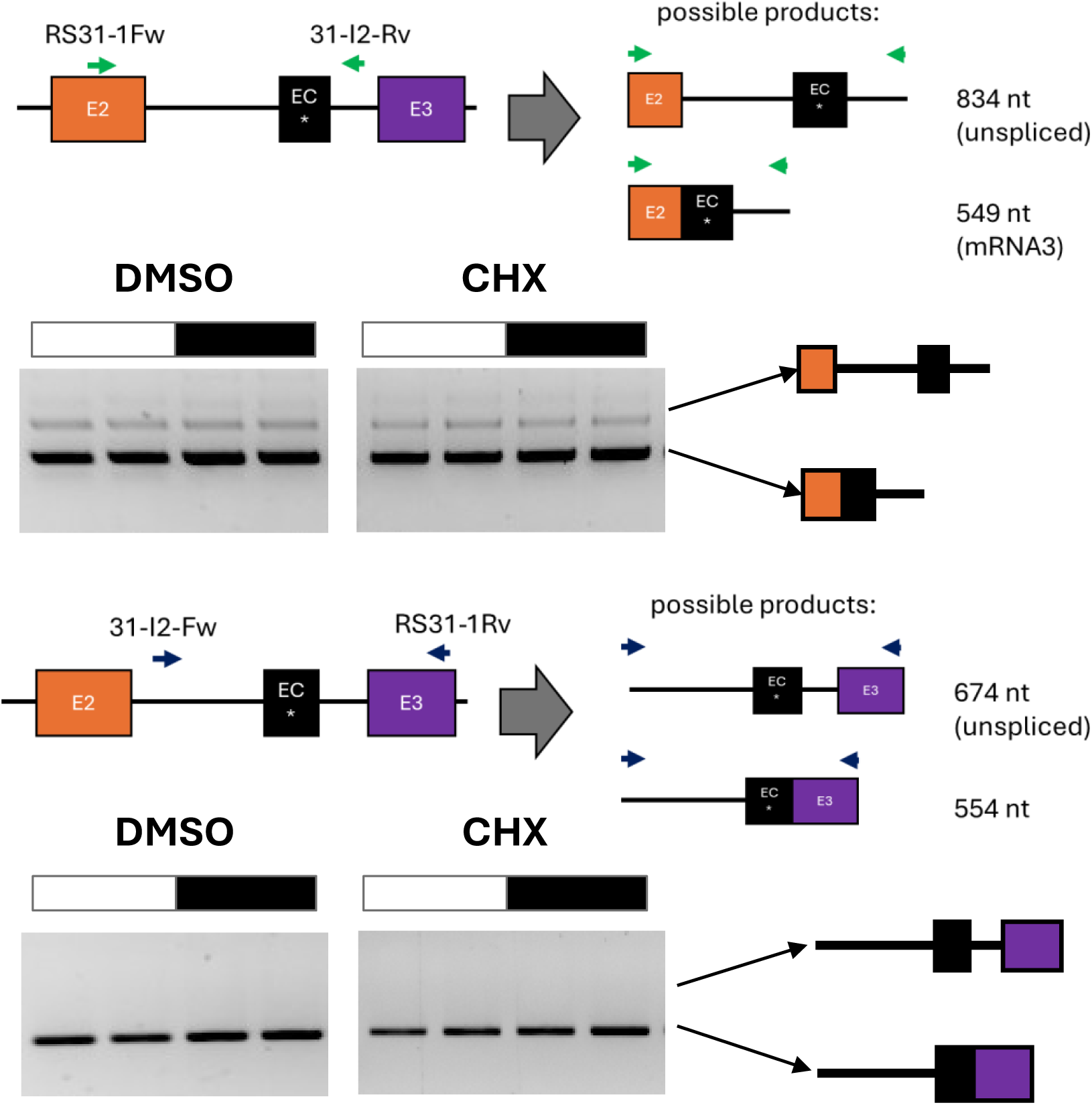

**Suppl. Figure S3.**
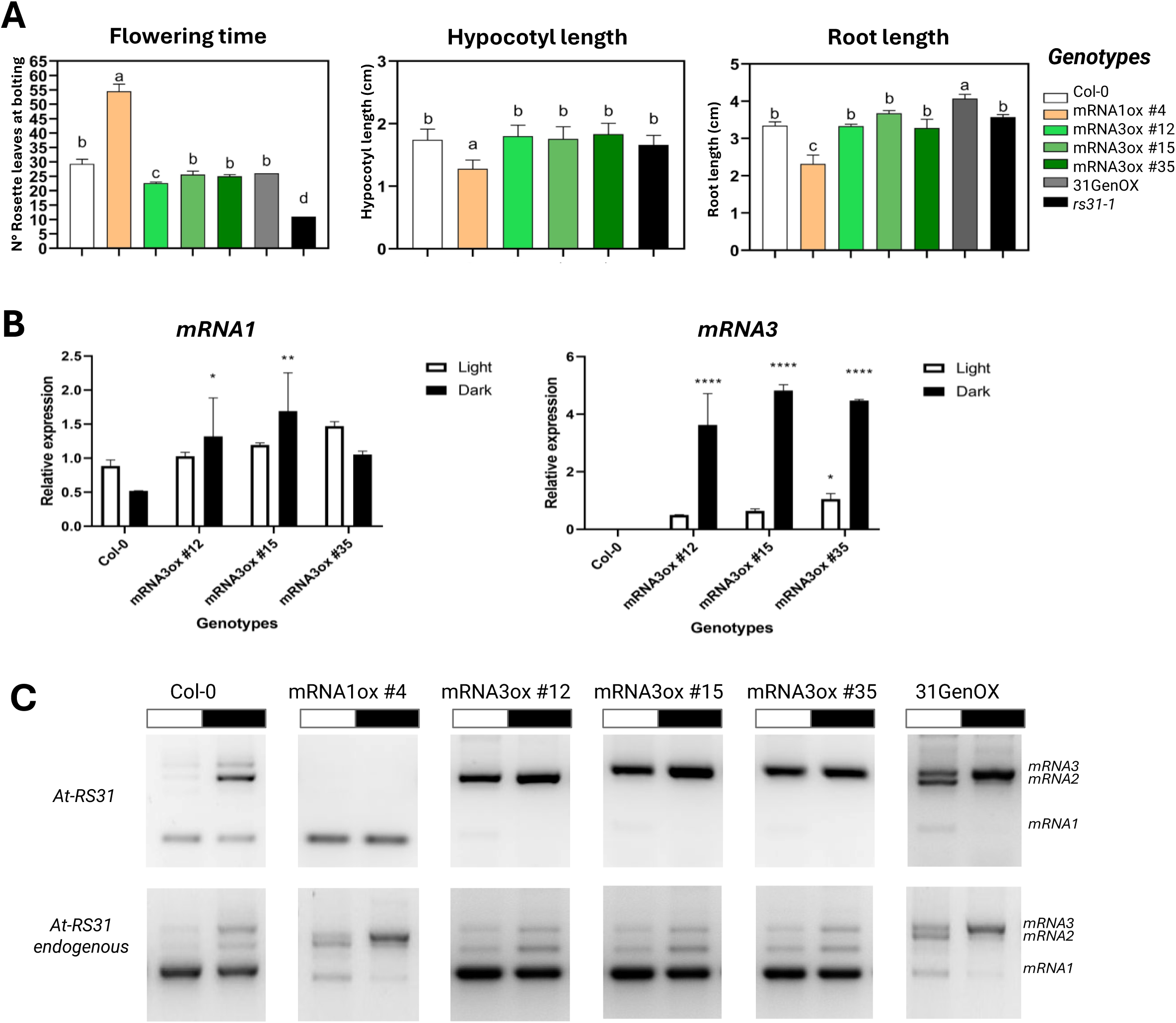

**Suppl. Figure S4.**
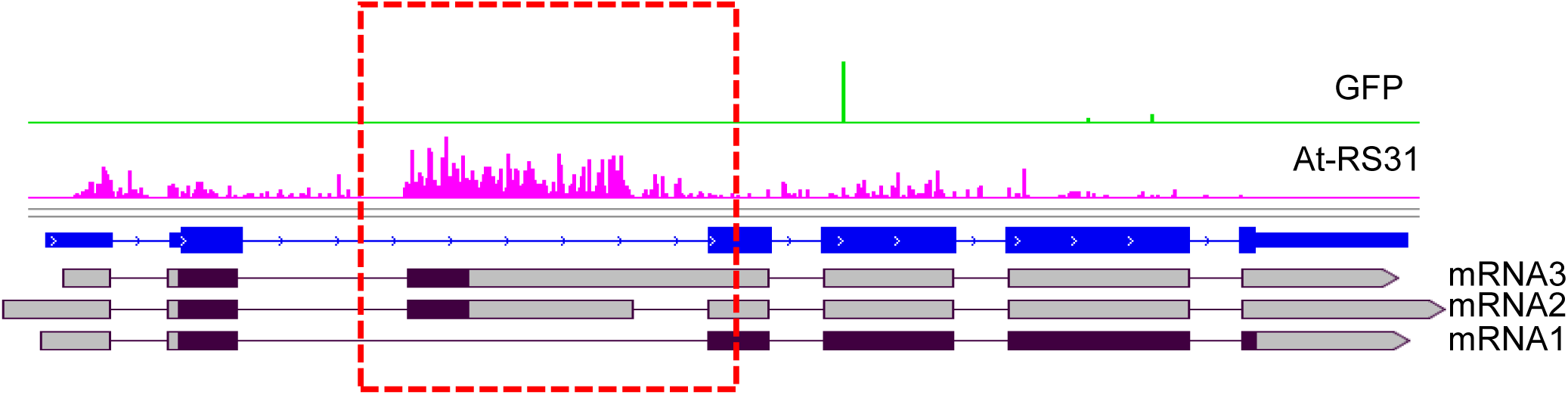

## References

1. Ashwal-Fluss R, Meyer M, Pamudurti NR, Ivanov A, Bartok O, Hanan M, Evantal N, Memczak S, Rajewsky N, Kadener S. 2014. circRNA biogenesis competes with pre-mRNA splicing. Molecular Cell 56, 55–66.

2. Barta A, Kalyna M, Reddy ASN. 2010. Implementing a rational and consistent nomenclature for serine/arginine-rich protein splicing factors (SR proteins) in plants. The Plant Cell 22, 2926–2929.

3. Benjamini Y, Hochberg Y. 1995. Controlling the False Discovery Rate: A Practical and Powerful Approach to Multiple Testing. Journal of the Royal Statistical Society. Series B (Methodological) 57, 289–300.

4. Berry S, Müller M, Rai A, Pelkmans L. 2022. Feedback from nuclear RNA on transcription promotes robust RNA concentration homeostasis in human cells. Cell Systems 13, 454–470.e15.

5. Bullard JH, Purdom E, Hansen KD, Dudoit S. 2010. Evaluation of statistical methods for normalization and differential expression in mRNA-Seq experiments. BMC Bioinformatics 11, 94.

6. Burgardt R, Bauer J, Reinhardt M, Rupp N, Engel C, Hellmann SL, Sack M, Weinberg Z, Wachter A. 2026. A structured RNA balances DEAD-box RNA helicase function in plant alternative splicing control. bioRxiv, DOI: 10.64898/2026.01.23.701338.

7. Chorostecki U, Bologna NG, Ariel F. 2023. The plant noncoding transcriptome: a versatile environmental sensor. The EMBO journal 42, e114400.

8. Clough SJ, Bent AF. 1998. Floral dip: a simplified method for Agrobacterium -mediated transformation of Arabidopsis thaliana. The Plant Journal 16, 735–743.

9. Conn VM, Hugouvieux V, Nayak A, et al. 2017. A circRNA from SEPALLATA3 regulates splicing of its cognate mRNA through R-loop formation. Nature Plants 3, 17053.

10. Fonouni-Farde C, Ariel F, Crespi M. 2021. Plant Long Noncoding RNAs: New Players in the Field of Post-Transcriptional Regulations. Non-Coding RNA 7, 12.

11. Fuchs A, Riegler S, Ayatollahi Z, et al. 2021. Targeting alternative splicing by RNAi: from the differential impact on splice variants to triggering artificial pre-mRNA splicing. Nucleic Acids Research 49, 1133–1151.

12. Gaggion N, Ariel F, Daric V, et al. 2021. ChronoRoot: High-throughput phenotyping by deep segmentation networks reveals novel temporal parameters of plant root system architecture. GigaScience 10(7):giab052.

13. Gaggion N, Boccardo NA, Bonazzola R, et al. 2026. ChronoRoot 2.0: An Open AI-Powered Platform for 2D Temporal Plant Phenotyping, GigaScience, giag018. DOI: 10.1093/gigascience/giag018

14. Godoy Herz MA, Kubaczka MG, Brzyżek G, Servi L, Krzyszton M, Simpson C, Brown J, Swiezewski S, Petrillo E, Kornblihtt AR. 2019. Light Regulates Plant Alternative Splicing through the Control of Transcriptional Elongation. Molecular Cell 73, 1066–1074.e3.

15. Göhring J, Jacak J, Barta A. 2014. Imaging of Endogenous Messenger RNA Splice Variants in Living Cells Reveals Nuclear Retention of Transcripts Inaccessible to Nonsense-Mediated Decay in Arabidopsis. The Plant Cell 26, 754–764.

16. Guo W, Tzioutziou N, Stephen G, Milne I, Calixto C, Waugh R, Brown JWS, Zhang R. 2019. A powerful and flexible tool for rapid and accurate differential expression and alternative splicing analysis of RNA-seq data for biologists. RNA Biol 18(11), 1574–1587.

17. Hartmann L, Wießner T, Wachter A. 2018. Subcellular Compartmentation of Alternatively Spliced Transcripts Defines SERINE/ARGININE-RICH PROTEIN30 Expression. Plant Physiology 176, 2886–2903.

18. Kalyna M, Lopato S, Voronin V, Barta A. 2006. Evolutionary conservation and regulation of particular alternative splicing events in plant SR proteins. Nucleic Acids Research 34, 4395–4405.

19. Kalyna M, Simpson CG, Syed NH, et al. 2012. Alternative splicing and nonsense-mediated decay modulate expression of important regulatory genes in Arabidopsis. Nucleic Acids Research 40, 2454–2469.

20. Kopp F, Mendell JT. 2018. Functional Classification and Experimental Dissection of Long Noncoding RNAs. Cell 172, 393–407.

21. Köster T, Venhuizen P, Lewinski M, et al. 2025. At-RS31 orchestrates hierarchical cross-regulation of splicing factors and integrates alternative splicing with TOR-ABA pathways. New Phytologist 247(2), 738–759.

22. Lampropoulos A, Sutikovic Z, Wenzl C, Maegele I, Lohmann JU, Forner J. 2013. GreenGate - A Novel, Versatile, and Efficient Cloning System for Plant Transgenesis (PJ Janssen, Ed.). PLoS ONE 8, e83043.

23. Law CW, Chen Y, Shi W, Smyth GK. 2014. voom: precision weights unlock linear model analysis tools for RNA-seq read counts. Genome Biology 15, R29.

24. Liu M, Wu S, Chen H, Wu S. 2012. Widespread translational control contributes to the regulation of Arabidopsis photomorphogenesis. Mol Syst Biol. 8, 566.

25. Marquez Y, Brown JWS, Simpson C, Barta A, Kalyna M. 2012. Transcriptome survey reveals increased complexity of the alternative splicing landscape in Arabidopsis. Genome Research 22, 1184–1195.

26. Mattick JS, Amaral PP, Carninci P, Carpenter S, Chang HY, Chen L, Chen R, Dean C, Dinger ME, Fitzgerald KA. 2023. Long non-coding RNAs: definitions, functions, challenges and recommendations. Nat Rev Mol Cell Biol 24(6), 430–447.

27. Meyer KD, Jaffrey SR. 2014. The dynamic epitranscriptome: N6-methyladenosine and gene expression control. Nature Reviews Molecular Cell Biology 15, 313–326.

28. Morton M, AlTamimi N, Butt H, Reddy ASN, Mahfouz M. 2019. Serine/Arginine-rich protein family of splicing regulators: New approaches to study splice isoform functions. Plant Science 283, 127–134.

29. Palusa SG, Ali GS, Reddy ASN. 2007. Alternative splicing of pre-mRNAs of Arabidopsis serine/arginine-rich proteins: regulation by hormones and stresses. Plant J 49(6), 1091–1107.

30. Palusa SG, Reddy ASN. 2010. Extensive coupling of alternative splicing of pre-mRNAs of serine/arginine (SR) genes with nonsense-mediated decay. New Phytol 2010 Jan;185(1):83–89.

31. Patro R, Duggal G, Love MI, Irizarry RA, Kingsford C. 2017. Salmon provides fast and bias-aware quantification of transcript expression. Nature Methods 14, 417–419.

32. Petrillo E. 2023. Don’t panic: An intron-centric guide to alternative splicing. The Plant Cell 35, 1752–1761.

33. Petrillo E, Godoy Herz MA, Fuchs A, et al. 2014. A Chloroplast Retrograde Signal Regulates Nuclear Alternative Splicing. Science 344, 427–430.

34. Riegler S, Servi L, Scarpin MR, et al. 2021. Light regulates alternative splicing outcomes via the TOR kinase pathway. Cell Reports 36(10), 109676.

35. Rigo R, Bazin J, Romero-Barrios N, et al. 2020. The Arabidopsis lnc RNA ASCO modulates the transcriptome through interaction with splicing factors. EMBO Reports 21(5), e48977.

36. Ritchie ME, Phipson B, Wu D, Hu Y, Law CW, Shi W, Smyth GK. 2015. Limma powers differential expression analyses for RNA-sequencing and microarray studies. Nucleic Acids Research 43(7):e47.

37. Rodríguez FS, Servi L, Quadrana L, Petrillo E. 2025. SuB3 a Simple Subcellular Fractionation Protocol for Localization Studies of Nucleic Acids and Proteins. bioRxiv DOI: 10.1101/2025.10.21.683662.

38. Romero-Barrios N, Legascue MF, Benhamed M, Ariel F, Crespi M. 2018. Splicing regulation by long noncoding RNAs. Nucleic Acids Research 46, 2169–2184.

39. Roundtree IA, Evans ME, Pan T, He C. 2017. Dynamic RNA Modifications in Gene Expression Regulation. Cell 169, 1187–1200.

40. Saha C, Saha S, Bhattacharyya NP. 2025. LncRNAOmics: A Comprehensive Review of Long Non-Coding RNAs in Plants. Genes 16(7), 765.

41. Saile J, Walter H, Denecke M, Lederer P, Schütz L, Hiltbrunner A, Lepp K, Lobato-Gil S, Beli P, Wachter A. 2025. A network of RS splicing regulatory proteins controls light-dependent splicing and seedling development. Plant Physiology 199(3), kiaf482.

42. Shi Q, Li H, Song J, Wu H, Zeng Y, Jiang J, Li S, Chen Z, He X. 2026. Human messenger RNA harbors widespread noncoding splice isoforms. Briefings in Bioinformatics 27, bbag068.

43. Soneson C, Love MI, Robinson MD. 2016. Differential analyses for RNA-seq: transcript-level estimates improve gene-level inferences. F1000Research 4, 1521.

44. Staiger D, Brown JWS. 2013. Alternative Splicing at the Intersection of Biological Timing, Development, and Stress Responses. The Plant Cell 25, 3640–3656.

45. Wagner RE, Frye M. 2021. Noncanonical functions of the serine-arginine-rich splicing factor (SR) family of proteins in development and disease. BioEssays 43.

46. Williamson L, Saponaro M, Boeing S, et al. 2017. UV Irradiation Induces a Non-coding RNA that Functionally Opposes the Protein Encoded by the Same Gene. Cell 168, 843–855.e13.

47. Xu S, Dong Q, Deng M, et al. 2021. The vernalization-induced long non-coding RNA VAS functions with the transcription factor TaRF2b to promote TaVRN1 expression for flowering in hexaploid wheat. Molecular Plant 14, 1525–1538.

48. Zhang R, Calixto CPG, Marquez Y, et al. 2017. A high quality Arabidopsis transcriptome for accurate transcript-level analysis of alternative splicing. Nucleic Acids Research 45, 5061–5073.

49. Zhang Y-C, He R-Q, Cheng Y, Wang D, Ariel F, Chen Y-Q. 2025. Long noncoding RNAs as molecular architects: Shaping plant functions and physiological plasticity. Molecular Plant 18, 1643–1671.

50. Zhang L, Zhang Y, Liu J, Li H, Liu B, Zhao T. 2023. N6-methyladenosine mRNA methylation is important for the light response in soybean. Frontiers in Plant Science 14, 1153840.

51. Zhao X, Li J, Lian B, Gu H, Li Y, Qi Y. 2018. Global identification of Arabidopsis lncRNAs reveals the regulation of MAF4 by a natural antisense RNA. Nature Communications 9, 5056.

52. Zheng X, Peng Q, Wang L, Zhang X, Huang L, Wang J, Qin Z. 2020. Serine/arginine-rich splicing factors: the bridge linking alternative splicing and cancer. International Journal of Biological Sciences 16, 2442–2453.

