## Supplementary material for "ALTERNATIVE SPLICING OF A CODING GENE PRODUCES A NUCLEAR REGULATORY LONG NON-CODING RNA": Suppl. Data S2

### Supplementary Data 2 -PTC containing isoforms in SR coding genes

| Gene name | Gene ID | Condtion Seedling Growth | Intron Upstream | Fully Spliced (coding) | Alt 3'ss or 5'ss | Intron retention | Intron Downstream | Freq. Non-coding | Total junctions splicing event | Minimum diff | Freq. Non-coding adjusted to introns in the vicinity | Including Possible IR | |
|---|---|---|---|---|---|---|---|---|---|---|---|---|---|
| SR30 | AT1G09140 | Light | 3474 | 1248 | 1460 | Not considered | 3233 | 0.54 | 2708 | 525 | 0.45 | 0.61 | |
| SR30 | AT1G09140 | Dark | 1501 | 332 | 1246 | Not considered | 1501 | 0.79 | 1578 | -77 | 0.83 | 0.78 | |
| SR34 | AT1G02840 | Light | 5048 | 5164 | 1272 | Not considered | 7180 | 0.20 | 6436 | 744 | 0.18 | 0.28 | |
| SR34 | AT1G02840 | Dark | 2439 | 1768 | 1223 | Not considered | 3231 | 0.41 | 2991 | 240 | 0.38 | 0.45 | |
| SR34a | AT3G49430 | Light | - | 2608 | 496 | Not considered | 3099 | 0.16 | 3104 | -5 | 0.16 | 0.16 | |
| SR34a | AT3G49430 | Dark | - | 262 | 408 | Not considered | 616 | 0.61 | 670 | -54 | 0.66 | 0.57 | |
| SR34b | AT4G02430 | Light | 507 | 186 | Not considered | 321 | 1008 | 0.63 | 507 | 0 | 0.63 | 0.63 | IR calculated from upstream intron junctions |
| SR34b | AT4G02430 | Dark | 2439 | 1768 | Not considered | 1223 | 3231 | 0.41 | 2991 | -552 | 0.50 | 0.28 | IR calculated from upstream intron junctions |
| RS31a | AT2G46610 | Light | 226 | 200 | 92 | 137 | 429 | 0.32 | 292 | 137 | 0.21 | 0.53 | IR calculated from upstream intron junctions |
| RS31a | AT2G46610 | Dark | 2439 | 1768 | 1223 | 240 | 3231 | 0.41 | 2991 | 240 | 0.38 | 0.45 | IR calculated from upstream intron junctions |
| RS31 | AT3G61860 | Light | 1104 | 1258 | 719 | Not considered | 2088 | 0.36 | 1977 | 111 | 0.34 | 0.40 | |
| RS31 | AT3G61860 | Dark | 689 | 401 | 808 | Not considered | 1000 | 0.67 | 1209 | -209 | 0.81 | 0.60 | |
| RS40 | AT4G25500 | Light | 1746 | 1353 | 471 | Not considered | 1964 | 0.26 | 1824 | 140 | 0.24 | 0.31 | |
| RS40 | AT4G25500 | Dark | 947 | 450 | 518 | Not considered | 1088 | 0.54 | 968 | 120 | 0.48 | 0.59 | |
| RS41 | AT5G52040 | Light | 3442 | 3052 | 477 | Not considered | 3682 | 0.14 | 3529 | 153 | 0.13 | 0.17 | |
| RS41 | AT5G52040 | Dark | 1142 | 802 | 340 | Not considered | 1277 | 0.30 | 1142 | 135 | 0.27 | 0.37 | |

SR30 AT1G09140

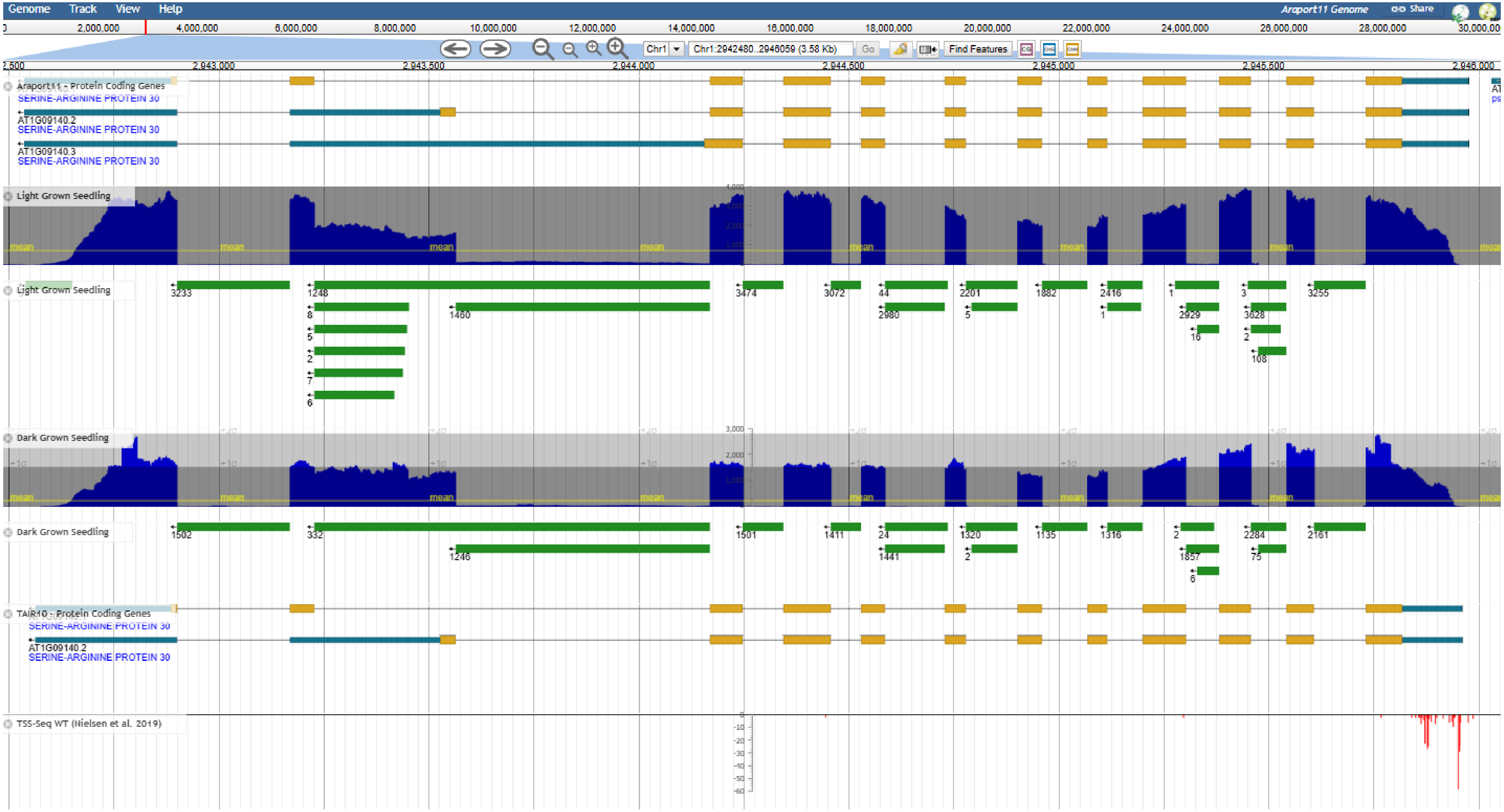

SR34 AT1G02840

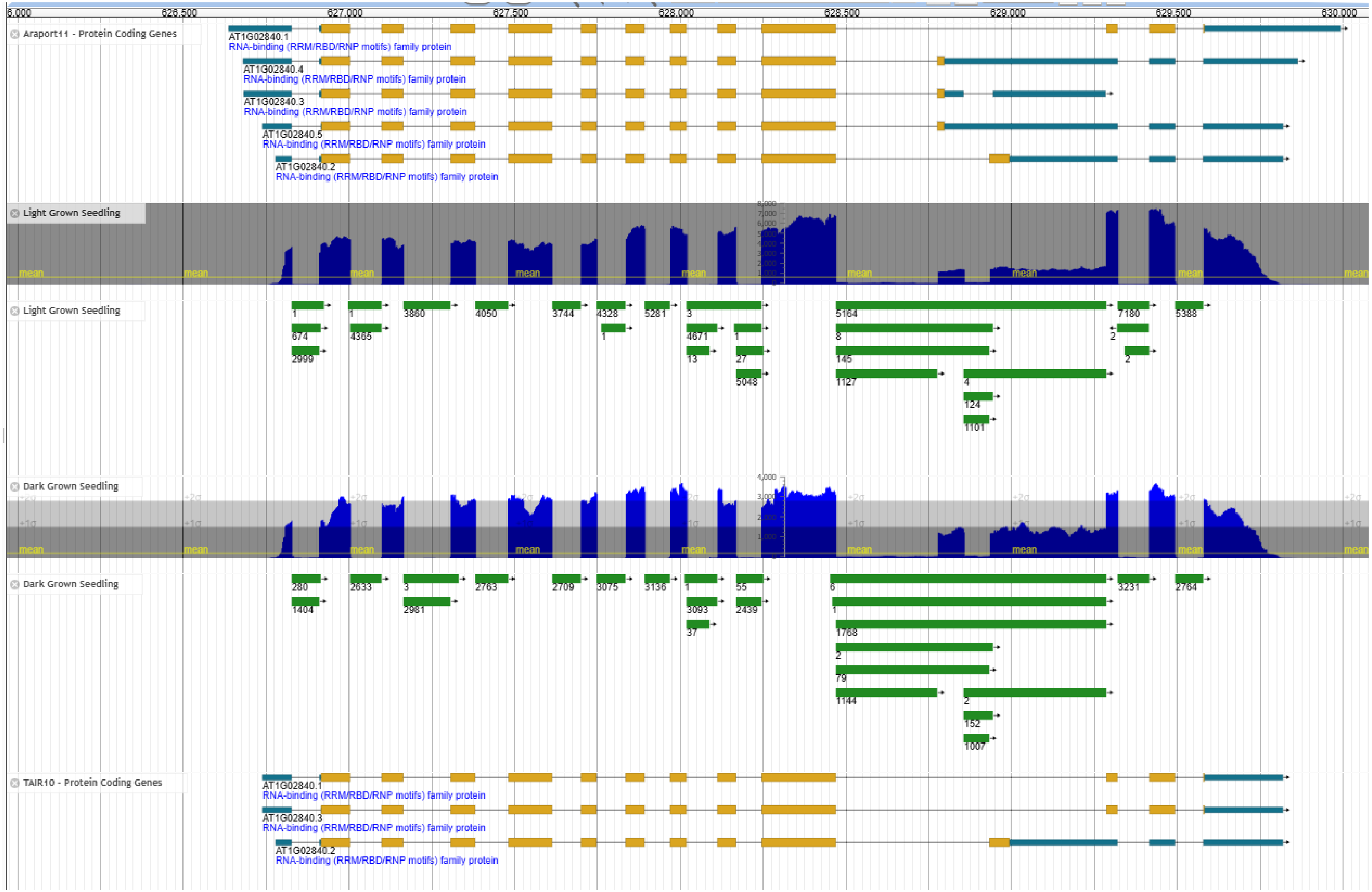

SR34a AT3G49430

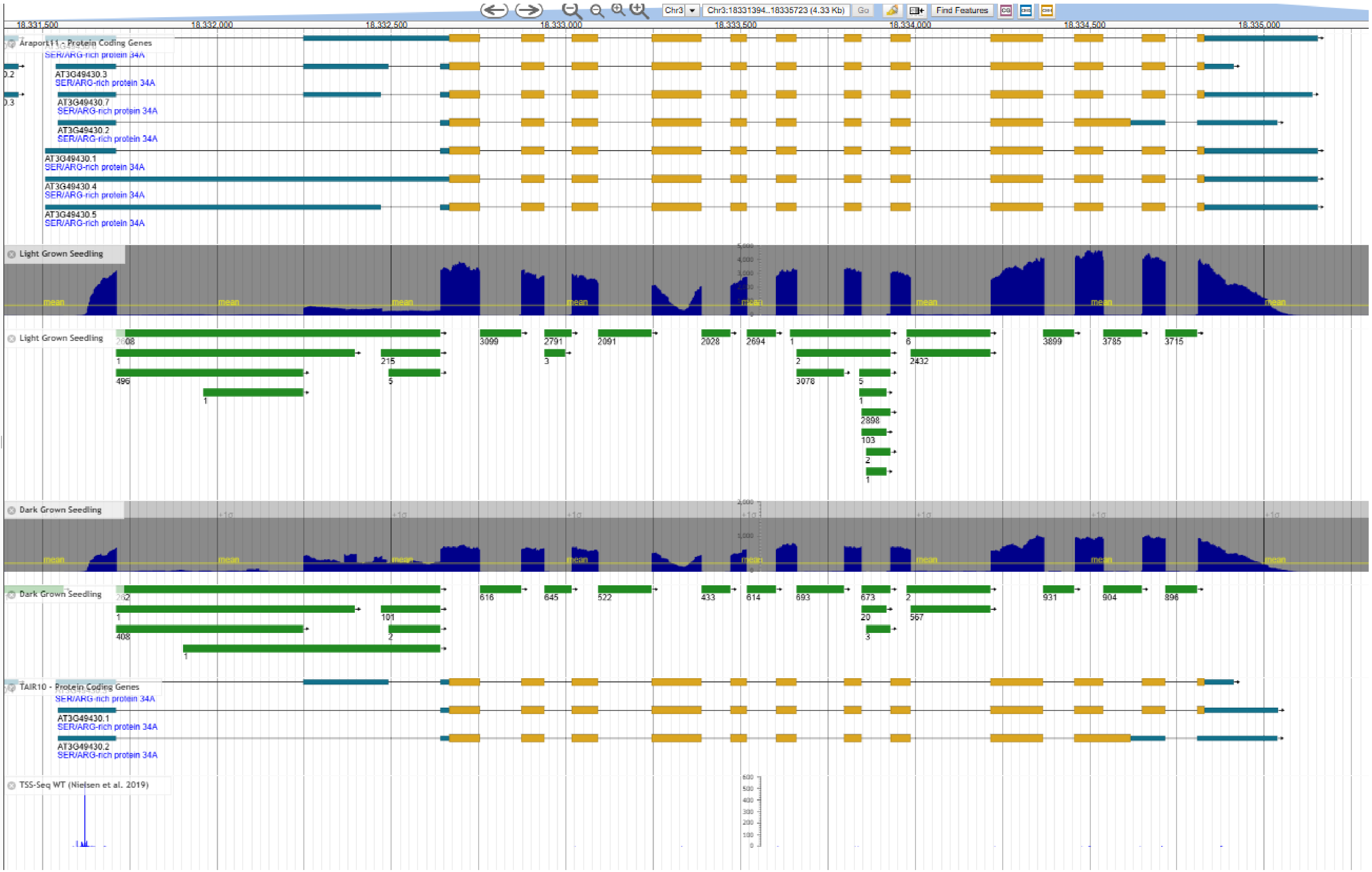

SR34b AT4G02430

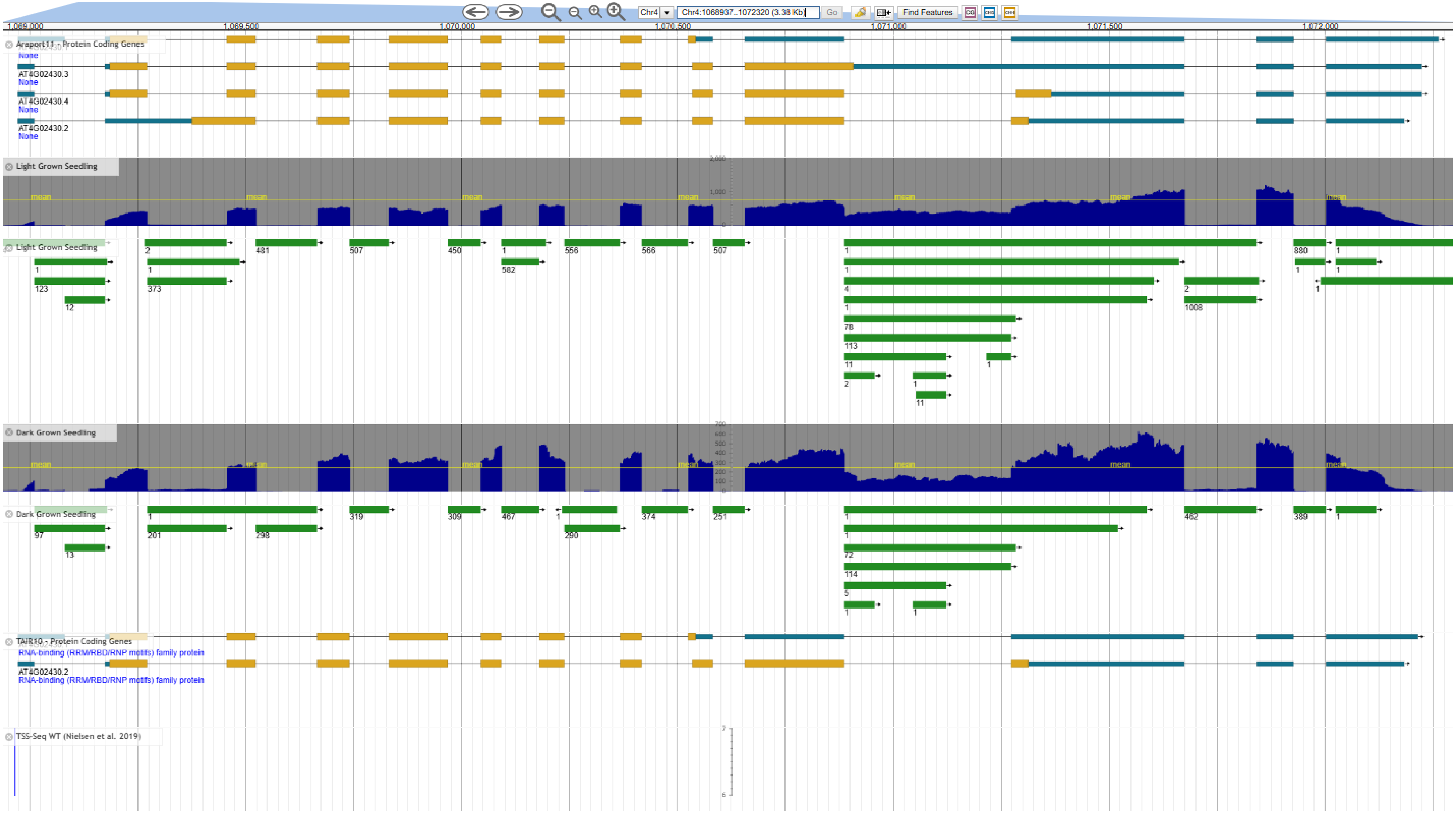

RS31a AT2G46610

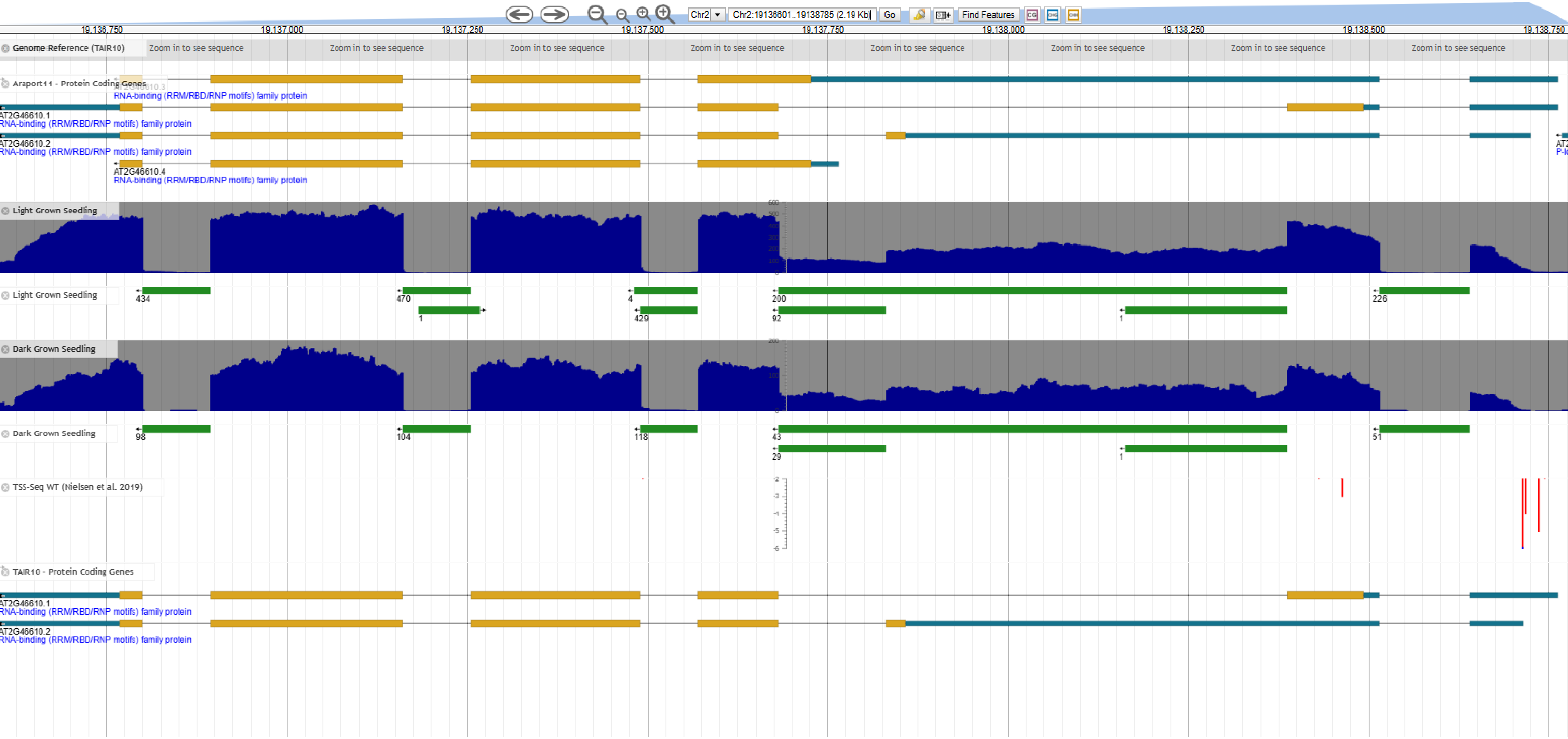

RS31 AT3G61860

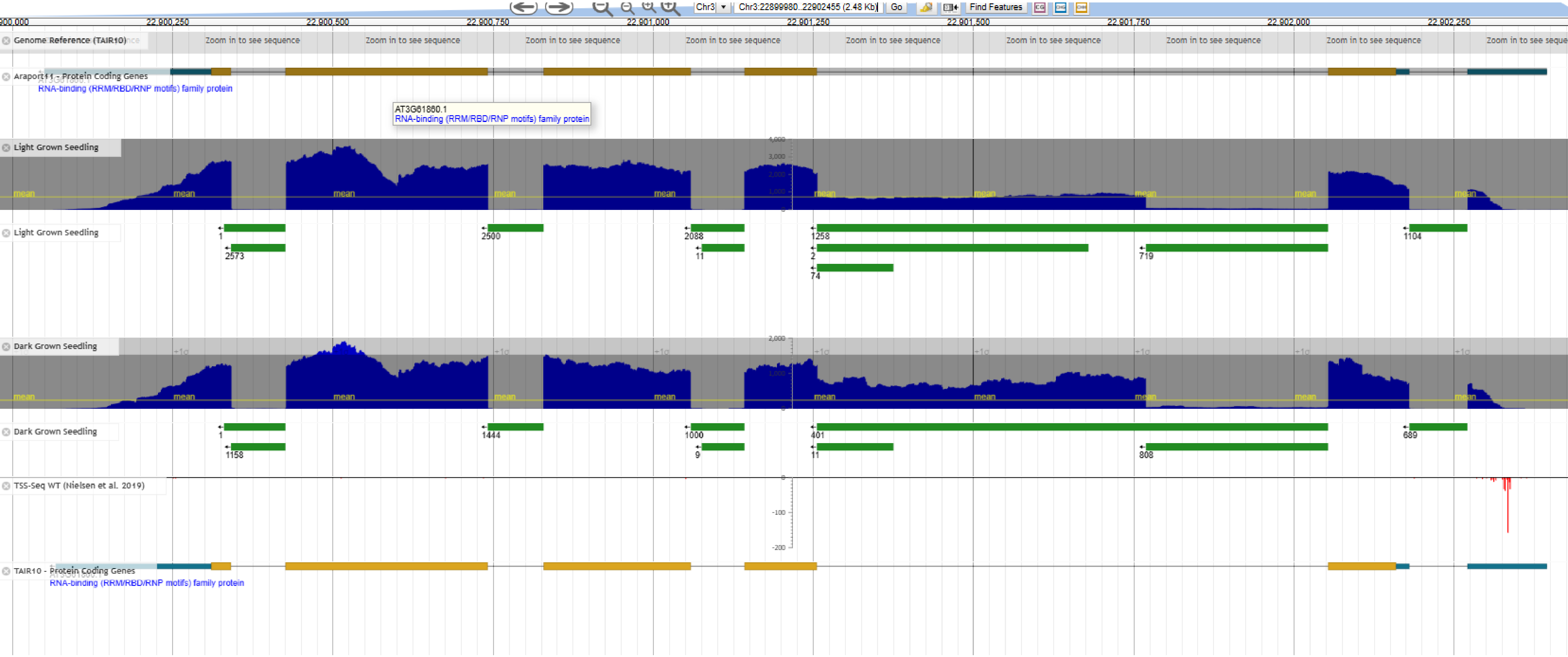

| | |
|---|---|
| RS40 | AT4G25500 |

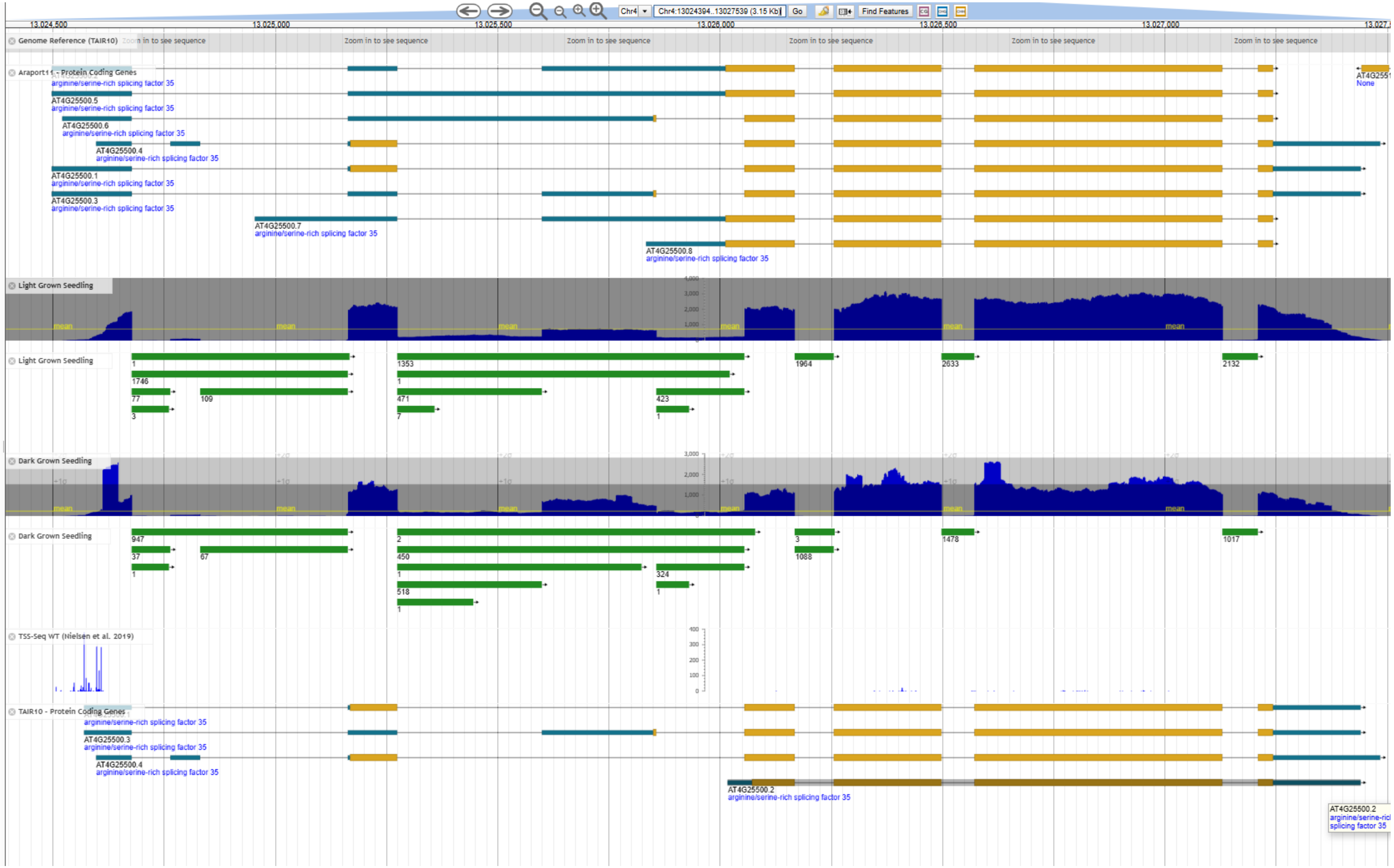

RS41 AT5G52040

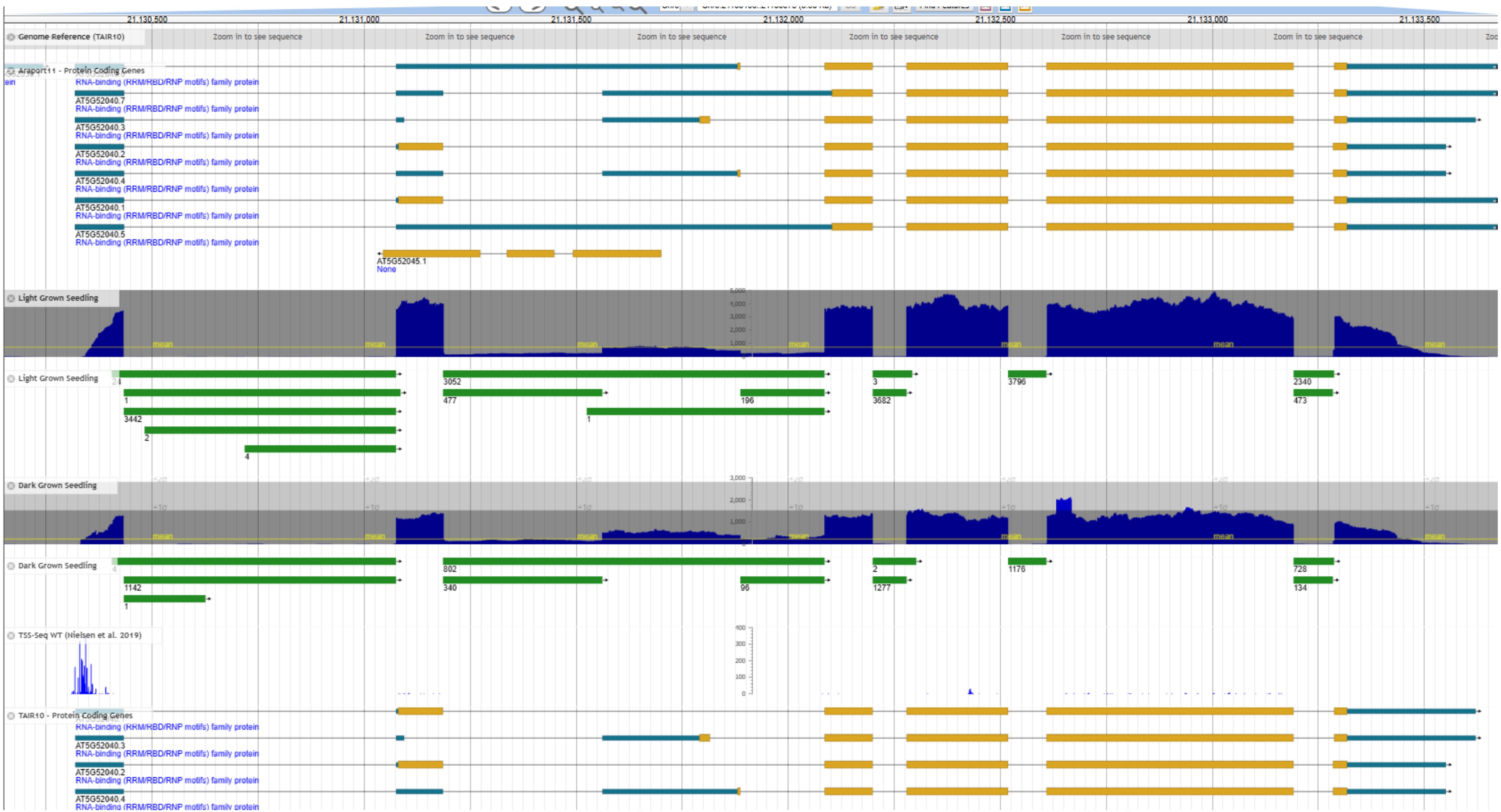

SR45 AT1G16610

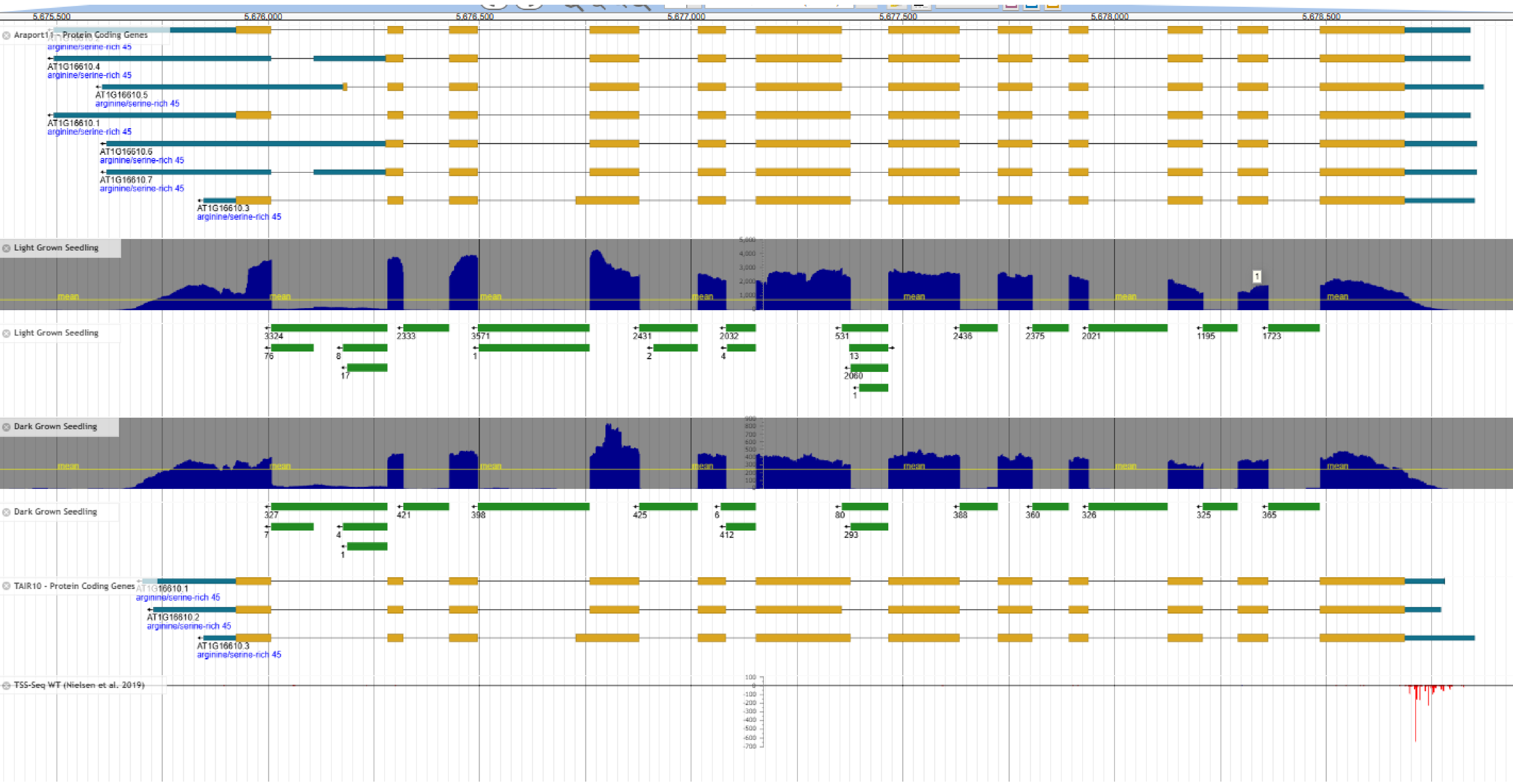

SR45a AT1G07350

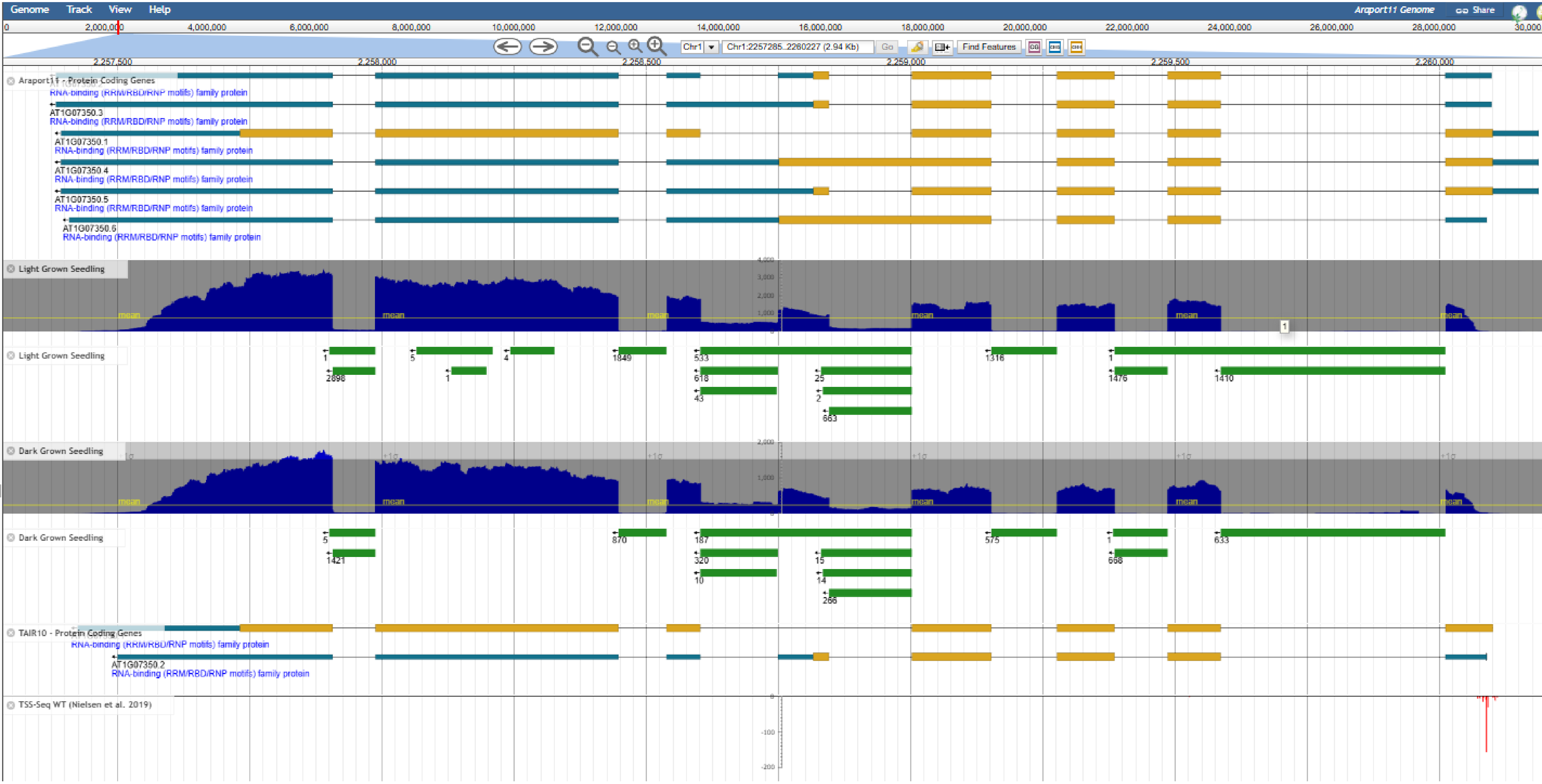

RSZ21 AT1G23860

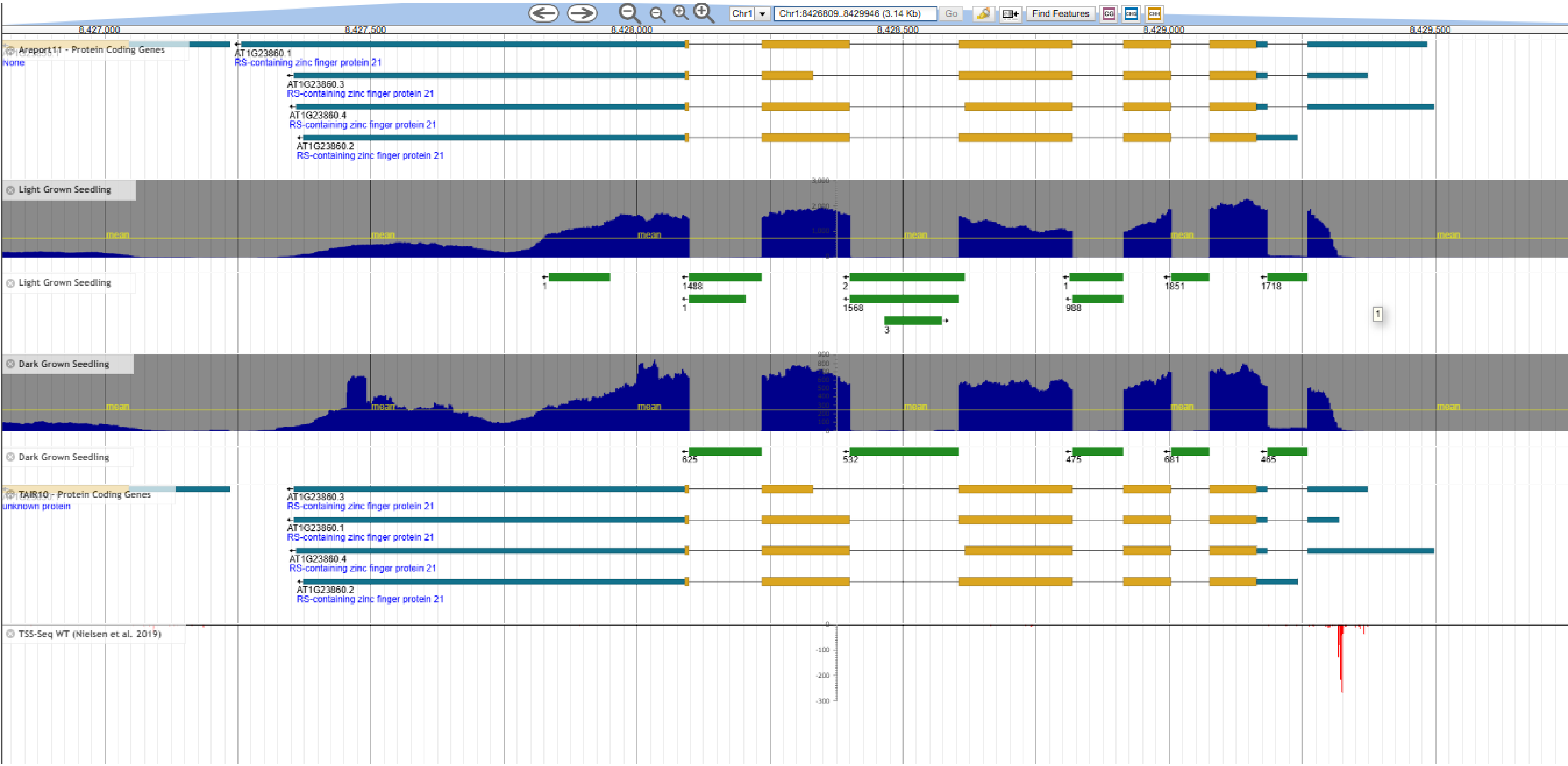

RSZ22 AT4G31580

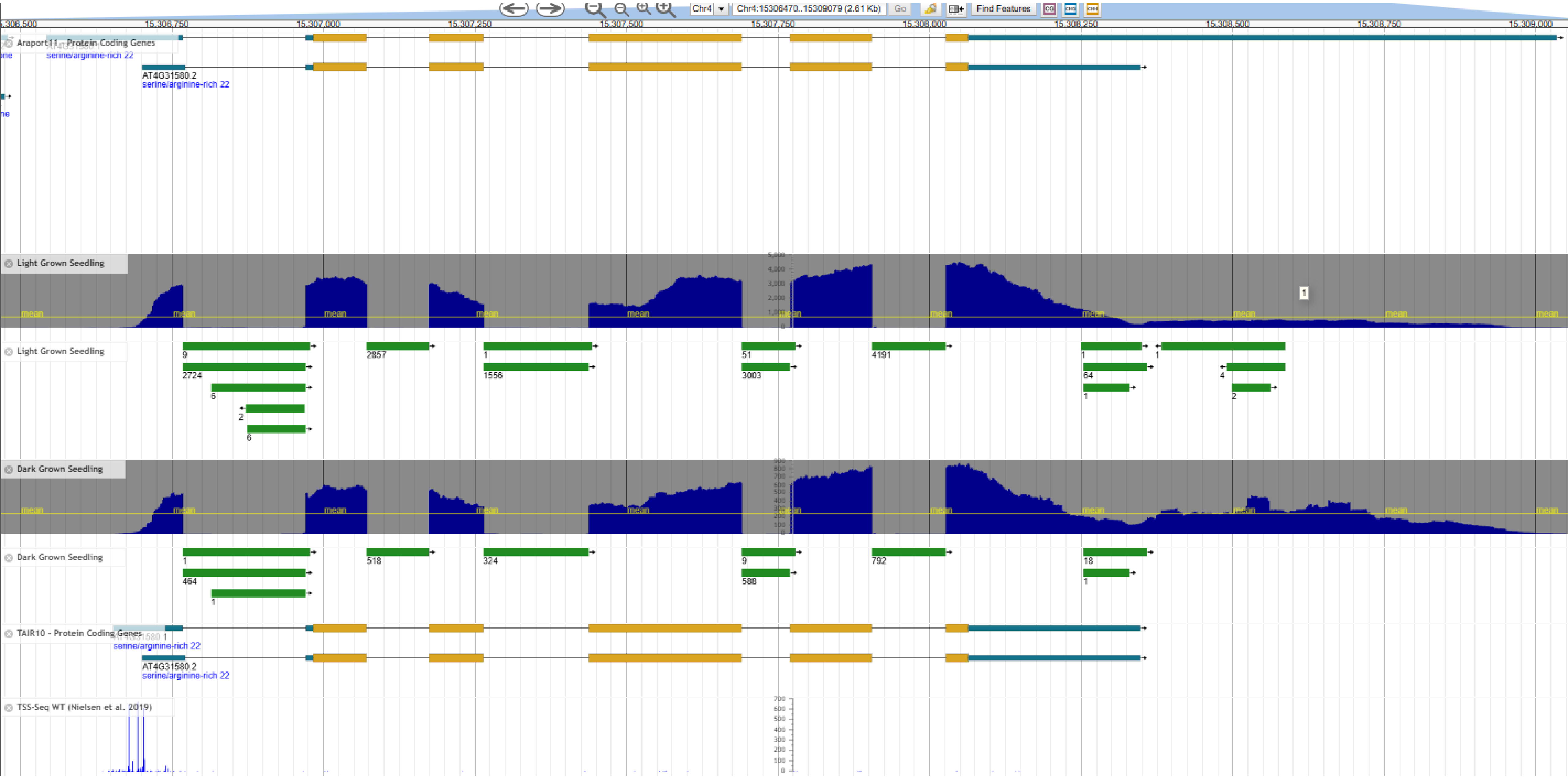

RSZ22a AT2G24590

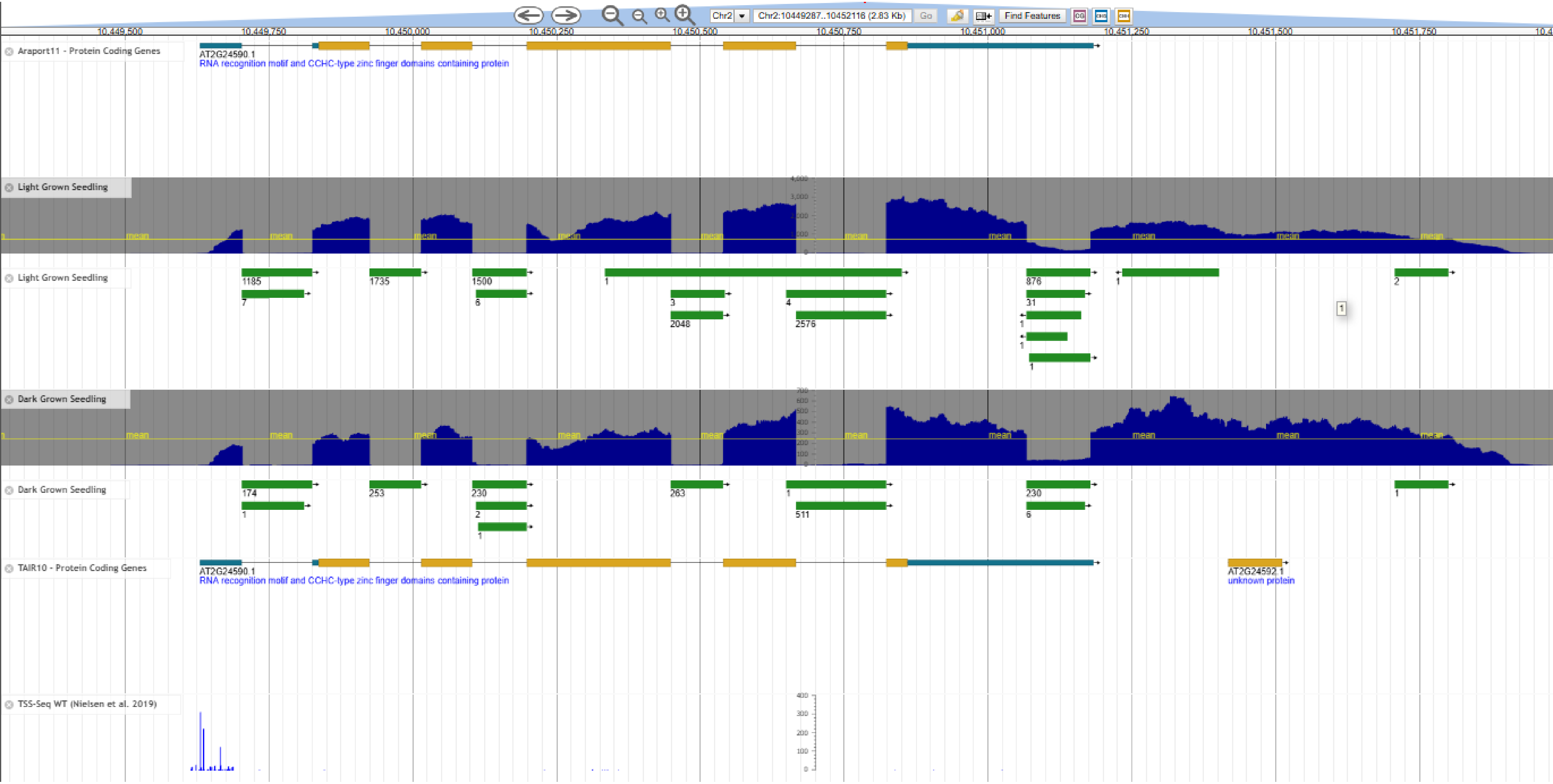

SC35 AT5G64200

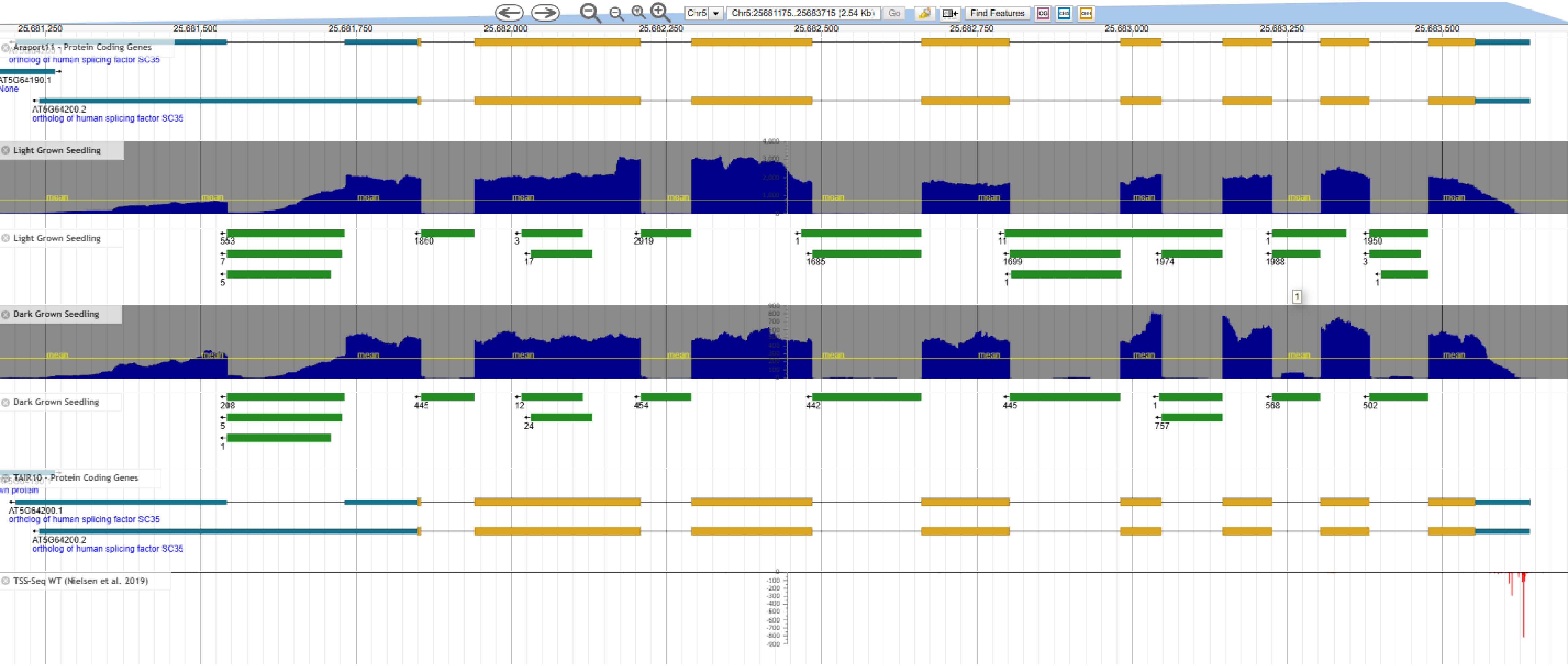

SCL28 AT5G18810

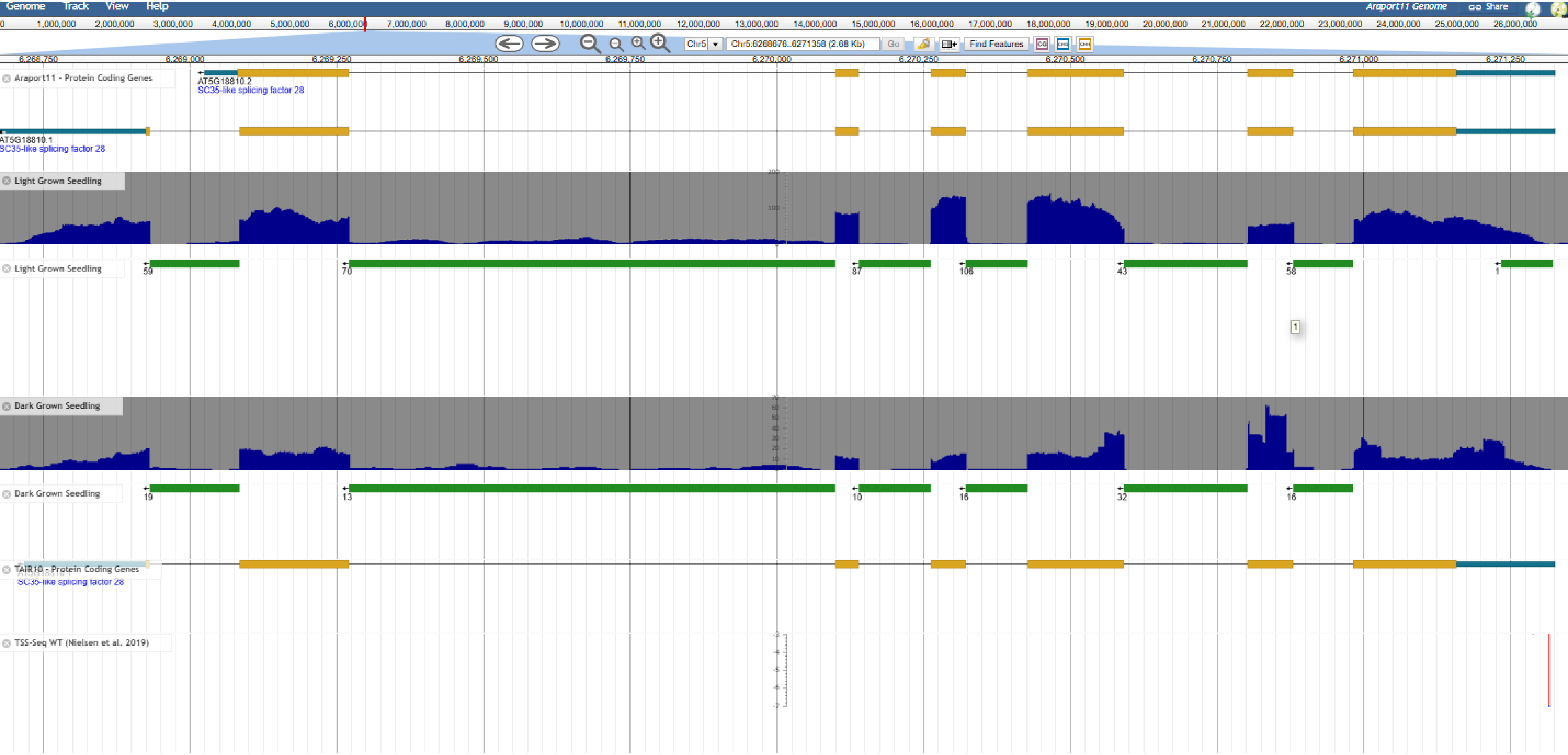

SCL30 AT3G55460

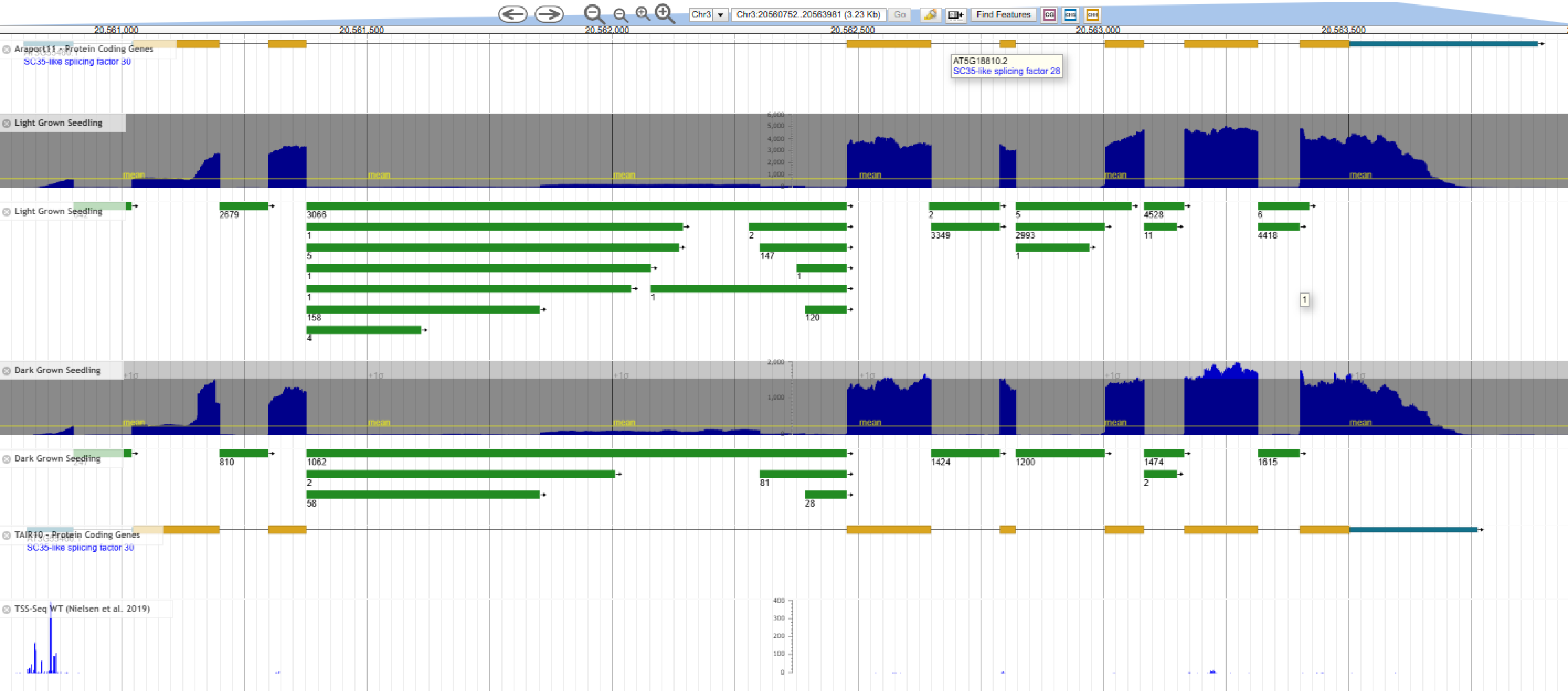

SCL30a AT3G13570

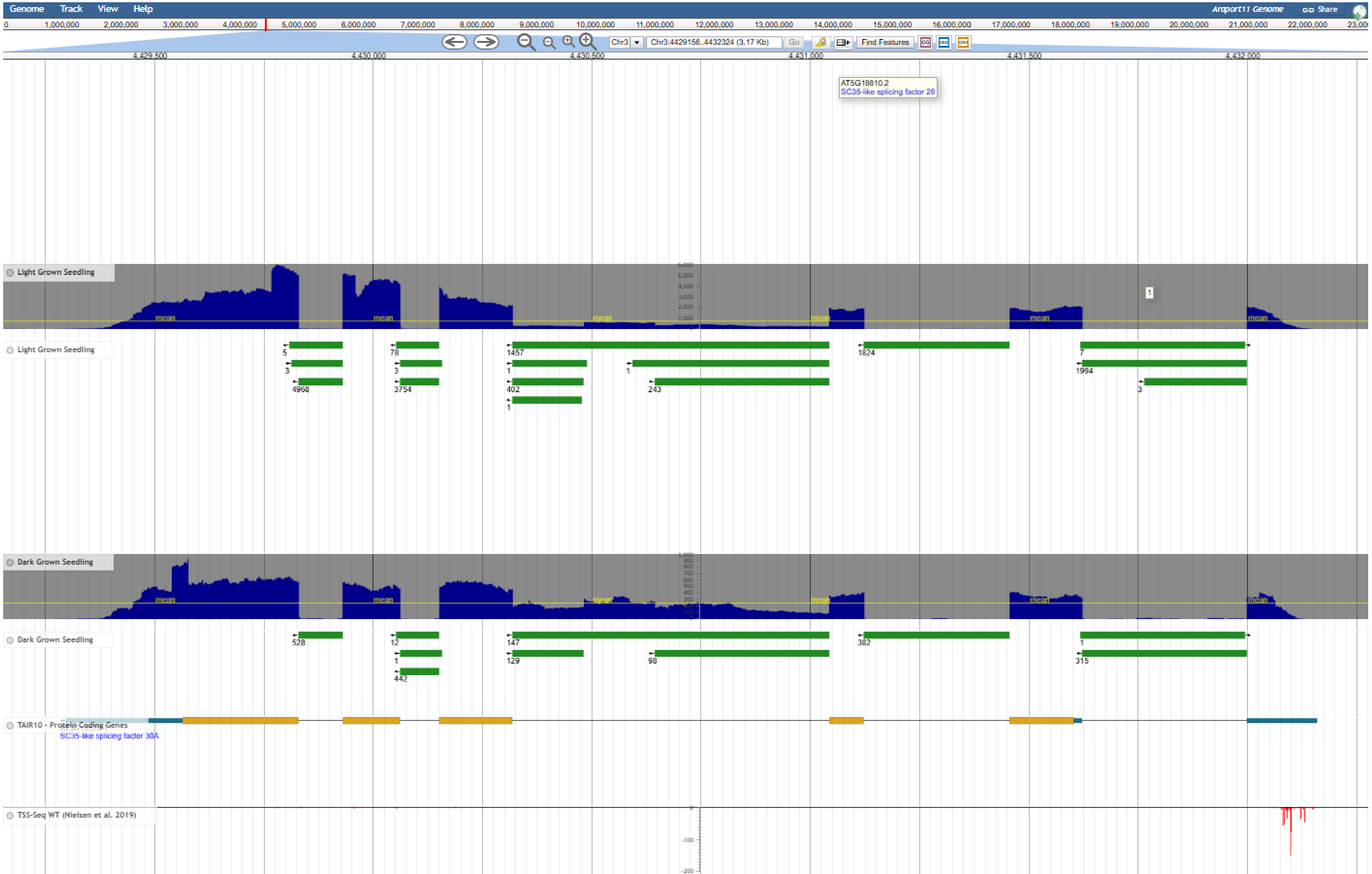

SCL33 AT1G55310

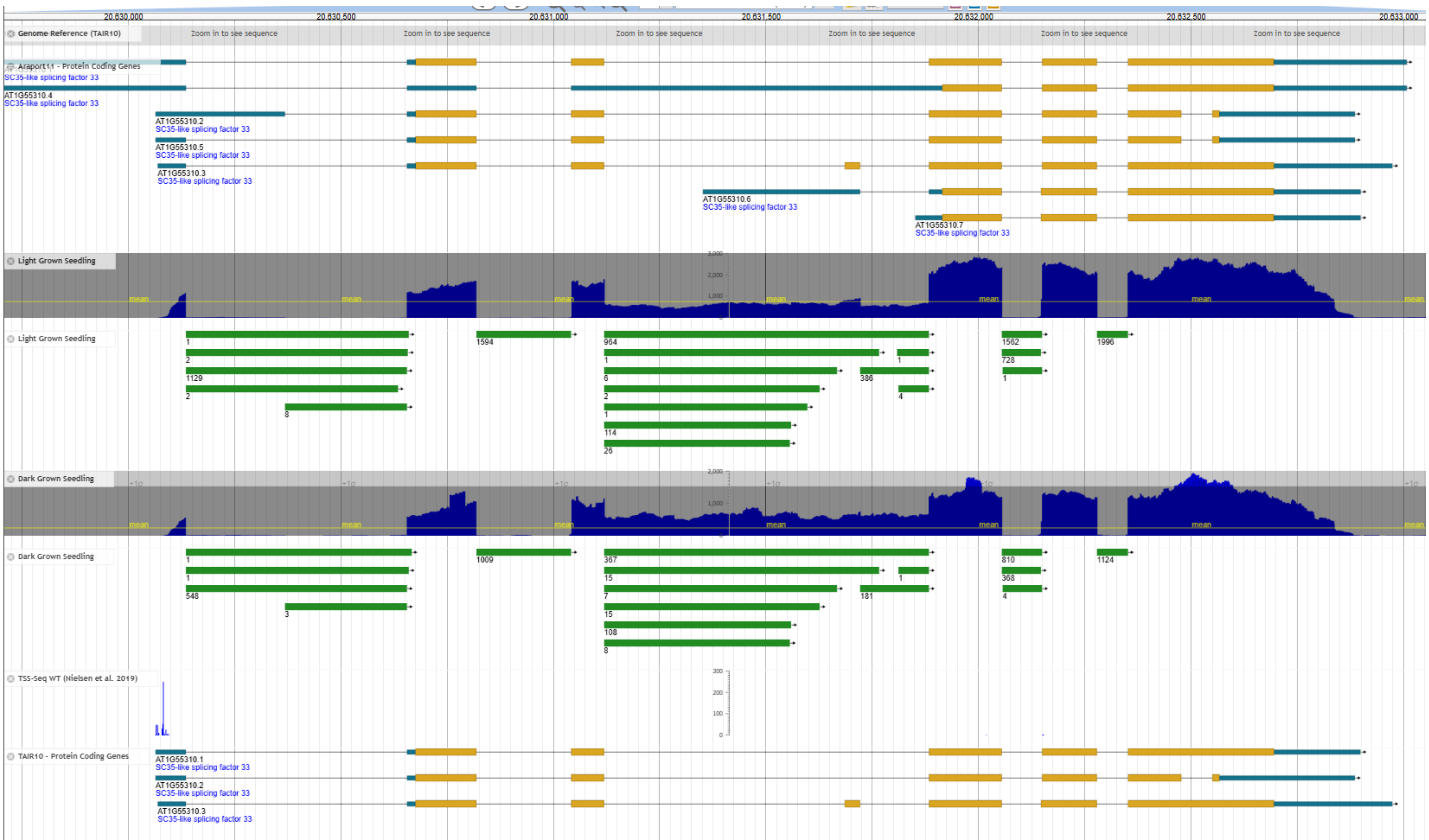

RS2Z32 AT3G53500

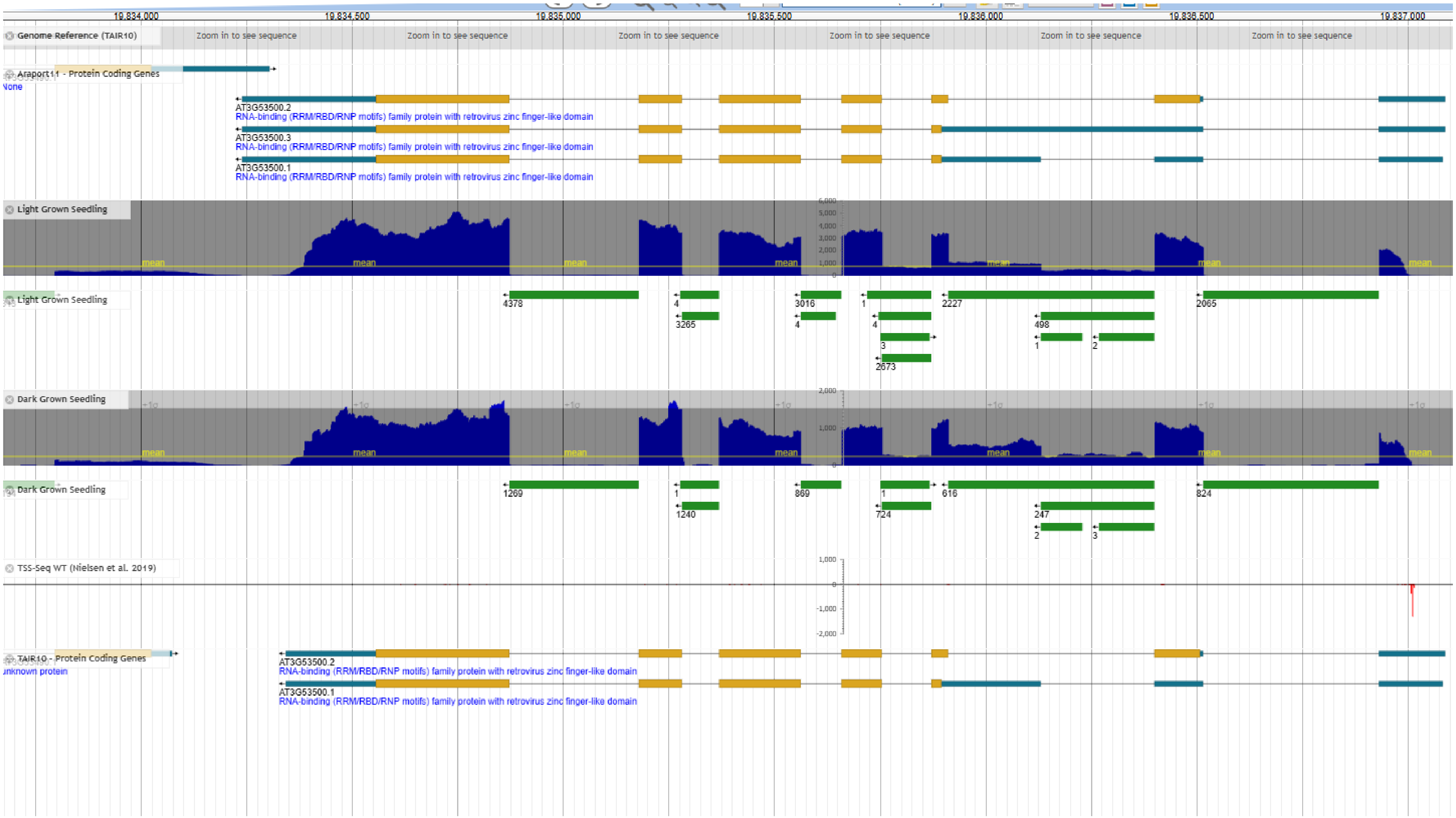

RS2Z33 AT2G37340

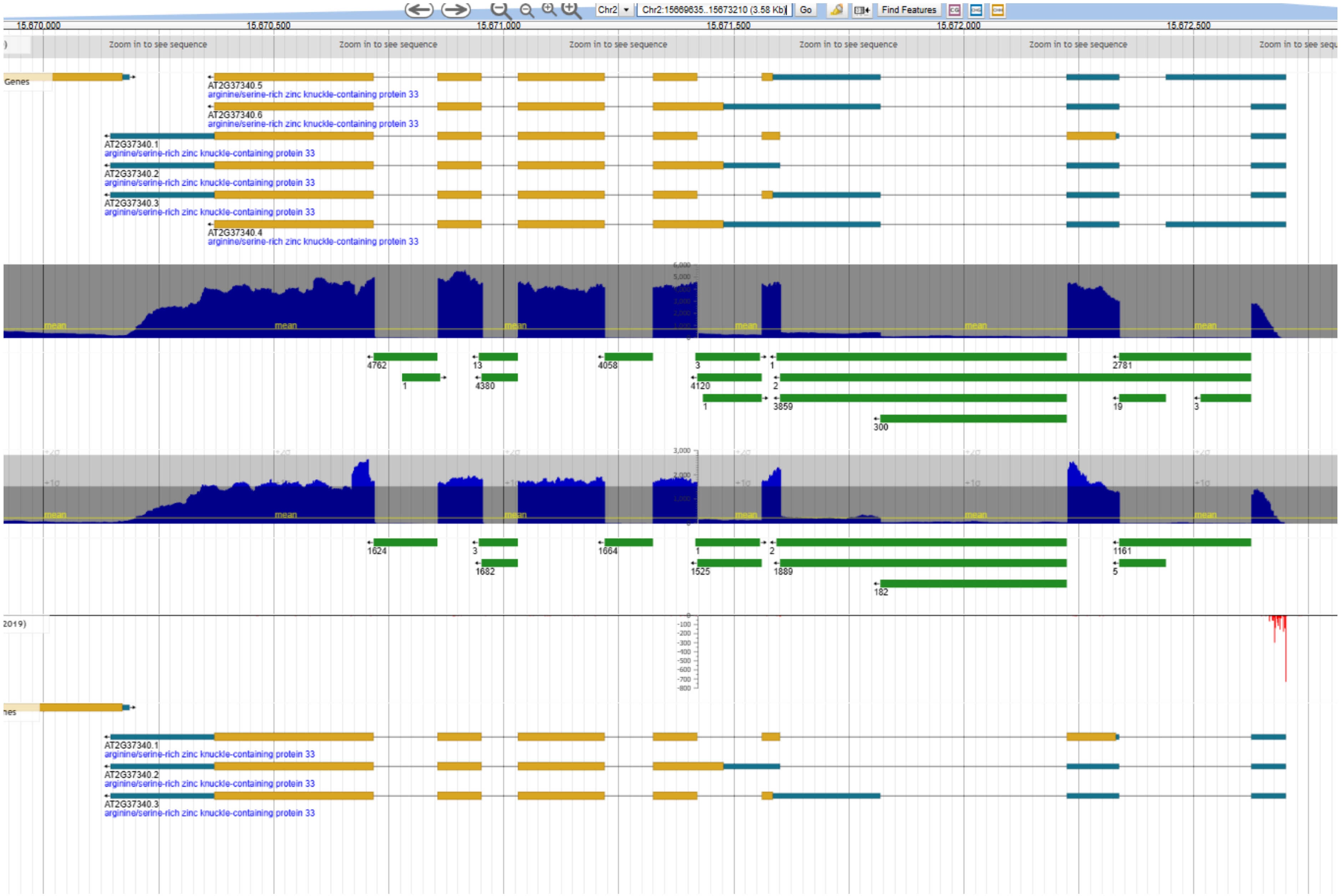
