## Supplementary material for "ALTERNATIVE SPLICING OF A CODING GENE PRODUCES A NUCLEAR REGULATORY LONG NON-CODING RNA": Suppl. Data S1

**Supplementary Data S1 – Rodriguez et al. 2026**

### Generation of splicing reporters

**At-RS31 region from exon 1 until exon 3 (5′ end only) with mutations to eliminate all possible PTCs:**

aaactcgagtgtacccatgatggtatgacgggctttttaagtcaaccaggctctgcttttctt
cgatgagcgtagtagtcgtcgtcgtcgtctagggtttgtttttcgtttcttctccgattgttca
gaggtaactttctccgataattatacttgctcatgtccatataccttattttatcccttacat
gttactaatttcgagttattggatcgttaggaattgcgaattaagataaagATGAGGcggccgC
CAGTGTTCGTCGGCAATTTCGAGTATGAAACTCGCCAGTCGGATCTGGAACGGTTGTTCGACAA
GTATGGGAGAGTCGACCGAGTGGACATGAAATCTGgtaaagtgtcttctaagttatacgccggt
gaaattttgttgtttgtccgtctcgaattcgaattcgtcttctggttgataaatcatctttgct
taggtttagtttctgcttgacctaggattctggatacttagaactagttccagtttcatacatg
atttgaaggtttatatatagtaccgttacatagaaaagattggttttattttcccgaatgtgt
taagtgtggtaggggtttatgcagtagagaggatggttgtattgagtgtttatttgttcgcag
gaacttttcttcaaaggaacaagatttgccatcttcttcttcttcctgcaaaatcatttctaca
ttttcatacctgcctttatcatcttgtgcctaccaatcttcactgactctagagaaactttgga
atttgtctgtctattaccaatcatcgatgcttcagatggaagtctattgaatccatcattgcca
tgccttgccttccatatctTG(ga)actcctccactcagaatgtttaagcatgcctggcaaatc
actcagattttG(t)aatataaaattggtgtttttaaaaataaaaatcctgaattcctgtgtt
atgctacaaatcttgtttttctactacatcatggaatgcctctacatctctcatattcaccagc
cttcacttatatgaaggtgttgtatatctatttagttattttcAT(ta)gggtaaacgagatat
gcagcaggttaatggacaggaggattgtcactctttgtgttgcgttttgAT(ta)atccttgtg
ctttcttttgttgcagGATATGggatccaaa

**The substrate for the mutations in *At-RS31* was ordered as a gBlock from IDT, with the following sequence to eliminate PTCs.**

cctgcctttatcatcttgtgcctaccaatcttcactactctagagaaactttggaatttgtctg
tctattaccaatcatcgatgcttcagatggaagtctattgaatccatcattgccatgccttgcc
ttccatatctTGactcctccactcagaatgtttaagcatgcctggcaaatcactcagattttG
aatataaaattggtgtttttaaaaataaaaatcctgaattcctgtgttatgctacaaatcttgt
tttctactacatcatggaatgcctctacatctctcatattcaccagccttcacttatatgaag
gtgttgtatatctatttagttattttcATgggtaaacgagatatgcagcaggttaatggacagg
aggattgtcactctttgtgttgcgttttgATatccttgtgctttcttttgttgcagGATATGgg
atccaaa

**The hygromycin cassette was amplified from pER8 using the following primers:**

jHyg-F1: aaaggatccCCTGAACTCACCGCGAC (BamHI)

jHyg-R1: tttcccgggCTATTCCTTTGCCCTCGG (XmaI)

**Generating:**

CTCTCAATCCAAATAATCTGCACCGGATCCCCTAGAATGAAAAAGCCTGAACTCACCGCGACGT
CTGTCGAGAAGTTTCTGATCGAAAAGTTCGACAGCGTCTCCGACCTGATGCAGCTCTCGGAGGG
CGAAGAATCTCGTGCTTTCAGCTTCGATGTAGGAGGGCGTGGATATGTCCTGCGGGTAAATAGC
TGCGCCGATGGTTTCTACAAAGATCGTTATGTTTATCGGCACTTTGCATCGGCCGCGCTCCCGA
TTCCGGAAGTGCTTGACATTGGGGAATTCAGCGAGAGCCTGACCTATTGCATCTCCCGCCGTGC

ACAGGGTGTCACGTTGCAAGACCTGCCTGAAACCGAACTGCCCGCTGTTCTGCAGCCGGTCGCG
GAGGCC*ATG*GATGCGATCGCTGCGGCCGATCTTAGCCAGACGAGCGGGTTCGGCCCATTCGGAC
CGCAAGGAATCGGTCAATACACTACATGGCGTGATTTCATATGCGCGATTGCTGATCCCCATGT
GTATCACTGGCAAACTGTG*ATG*GACGACACCGTCAGTGCGTCCGTCGCGCAGGCTCTCGATGAG
CTG*ATG*CTTTGGGCCGAGGACTGCCCCGAAGTCCGGCACCTCGTGCACGCGGATTTCGGCTCCA
ACAATGTCCTGACGGACAATGGCCGCATAACAGCGGTCATTGACTGGAGCGAGGCGATGTTCGG
GGATTCCCAATACGAGGTCGCCAACATCTTCTTCTGGAGGCCGTGGTTGGCTTGTATGGAGCAG
CAGACGCGCTACTTCGAGCGGAGGCATCCGGAGCTTGCAGGATCGCCGCGGCTCCGGGCGTATA
TGCTCCGCATTGGTCTTGACCAACTCTATCAGAGCTTGGTTGACGGCAATTTCGATGATGCAGC
TTGGGCGCAGGGTCGATGCGACGCAATCGTCCGATCCGGAGCCGGGACTGTCGGGCGTACACAA
ATCGCCCGCAGAAGCGCGGCCGTCTGGACCGATGGCTGTGTAGAAGTACTCGCCGATAGTGGAA
ACCGACGCCCCAGCACTCGTCCGAGGGCAAAGGAATAGCGATCGTTCAAACATTTGGCAATAAA
GTTTCTTAAGATTGAATCCTGTTGCCGGTCTT

**~1030 bps**

Using cDNA (*mRNA3*) or genomic DNA (*At-RS31*) as templates for sequential PCRs, together with the synthesized fragment of *At-RS31* and the hygromycin cassette, we were able to generate the following reporter constructs:

**RS31-jHyg**

aaactcgag**tgtacccatgatggtatgacgggctttttaagtcaaccaggctctgcttttctt**
**cgatgagcgtagtagtcgtcgtcgtcgtctagggtttgtttttcgtttcttctccgattgttca**
**gag**gtaacttttctccgataattatacttgctcatgtccatataccttattttatcccttacat
gttactaatttcgagttattggatcgttag**gaattgcgaattaagataaag**ATGAGGcggccgC
CAGTGTTCGTCGGCAATTTCGAGTATGAAACTCGCCAGTCGGATCTGGAA***CGGTTGTTCGACAA***
***GTATG***GGAGAGTCGACCGAGTGGACATGAAATCTGgtaaagtgtcttctaagttatacgccggt
gaaatttgttgtttgtccgtctcgaattcgaattcgtcttctggttgataaatcatctttgct
taggtttagtttctgcttgacctaggattctggatacttagaactagttccagtttcatacatg
atttgaaggtttatatatagtaccgttacatagaaaagattggtttatttttcccgaatgtgt
taagtgtggtagggggtttatgcagtagagaggatggttgtattgagtgtttatttgttcgcag
**gaacttttcttcaaaggaacaagatttgccatcttcttcttcttcctgcaaatcatttctaca**
**ttttcatacctgcctttatcatcttgtgcctaccaatcttcactactctagagaaactttggaa**
**tttgtctgtctattaccaatcatcgatgcttcagatggaagtctattgaatccatcattgccat**
**gccttgccttccatatctTGactcctccactcagaatgtttaagcatgcctggcaaatcactca**
**gattttGaatataaaattggtgttttaaaaataaaaatcctgaattcctgtgttatgctaca**
**aatcttgtttttctactacatcatggaatgcctctacatctctcatattcaccagccttcactt**
**atatgaag*gtgttgtatatctatttagttattttcATgggtaaacgagatatgcagcaggttaa***
***tggacaggaggattgtcactctttgtgttgcgttttgATatccttgtgctttcttttgttgcag***
GATATG**GGATCCCCTGAACTCACCGCGACGT*CTGTCGAGAAGTTTCTGATC*GAAAAGTTCGACA**
**GCGTCTCCGACCTGATGCAGCTCTCGGAGGGCGAAGAATCTCGTGCTTTCAGCTTCGATGTAGG**
**AGGGCGTGGATATGTCCTGCGGGTAAATAGCTGCGCCGATGGTTTCTACAAAGATCGTTATGTT**
**TATCGGCACTTTGCATCGGCCGCGCTCCCGATTCCGGAAGTGCTTGACATTGGGGCATTCAGCG**
**AGAGCCTGACCTATTGCATCTCCCGCCGTGCACAGGGTGTCACGTTGCAAGACCTGCCTGAAAC**
**CGAACTGCCCGCTGTTCTGCAGCCGGTCGCGGAGGCCATGGATGCGATCGCTGCGGCCGATCTT**
**AGCCAGACGAGCGGGTTCGGCCCATTCGGACCGCAAGGAATCGGTCAATACACTACATGGCGTG**
**ATTTCATATGCGCGATTGCTGATCCCCATGTGTATCACTGGCAAACTGTGATGGACGACACCGT**
**CAGTGCGTCCGTCGCGCAGGCTCTCGATGAGCTGATGCTTTGGGCCGAGGACTGCCCCGAAGTC**
**CGGCACCTCGTGCACGCGGATTTCGGCTCCAACAATGTCCTGACGGACAATGGCCGCATAACAG**
**CGGTCATTGACTGGAGCGAGGCGATGTTCGGGGATTCCCAATACGAGGTCGCCAACATCTTCTT**

CTGGAGGCCGTGGTTGGCTTGTATGGAGCAGCAGACGCGCTACTTCGAGCGGAGGCATCCGGAG
CTTGCAGGATCGCCGCGGCTCCGGGCGTATATGCTCCGCATTGGTCTTGACCAACTCTATCAGA
GCTTGGTTGACGGCAATTTCGATGATGCAGCTTGGGCGCAGGGTCGATGCGACGCAATCGTCCG
ATCCGGAGCCGGGACTGTCGGGCGTACACAAATCGCCCGCAGAAGCGCGGCCGTCTGGACCGAT
GGCTGTGTAGAAGTACTCGCCGATAGTGGAAACCGACGCCCCAGCACTCGTCCGAGGGCAAAGG
AATAGCCCGGGAAAA

**mRNA3-jHyg**

aaactcgagtgtacccatgatggtatgacgggctttttaagtcaaccaggctctgcttttctt
cgatgagcgtagtagtcgtcgtcgtctagggtttgtttttcgtttcttctccgattgttca
gaggtaacttttctccgataattatacttgctcatgtccatataccttattttatcccttacat
gttactaatttcgagttattggatcgttaggaattgcgaattaagataaagATGAGGcggccgC
CAGTGTTCGTCGGCAATTTCGAGTATGAAACTCGCCAGTCGGATCTGGAACGGTTGTTCGACAA
GTATGGGAGAGTCGACCGAGTGGACATGAAATCTGgaacttttcttcaaaggaacaagatttgc
catcttcttcttcttcctgcaaaatcatttctacattttcatacctgcctttatcatcttgtgc
ctaccaatcttcactactctagagaaactttggaatttgtctgtctattaccaatcatcgatgc
ttcagatggaagtctattgaatccatcattgccatgccttgccttccatatctgTactcctcca
ctcagaatgtttaagcatgcctggcaaatcactcagattttGaatataaaattgggtgtttta
aaaataaaaatcctgaattcctgtgttatgctacaaatcttgtttttctactacatcatggaat
gcctctacatctctcatattcaccagccttcacttatatgaag*gtgttgtatatctatttagtt*
*attttcATgggtaaacgagatatgcagcaggttaatggacaggaggattgtcactctttgtgtt*
*gcgttttgATatccttgtgctttcttttgttgcag*GATATGGGATCCCCTGAACTCACCGCGAC
GTCTGTCGAGAAGTTTCTGATCGAAAAGTTCGACAGCGTCTCCGACCTGATGCAGCTCTCGGAG
GGCGAAGAATCTCGTGCTTTCAGCTTCGATGTAGGAGGGCGTGGATATGTCCTGCGGGTAAATA
GCTGCGCCGATGGTTTCTACAAAGATCGTTATGTTTATCGGCACTTTGCATCGGCCGCGCTCCC
GATTCCGGAAGTGCTTGACATTGGGGCATTCAGCGAGAGCCTGACCTATTGCATCTCCCGCCGT
GCACAGGGTGTCACGTTGCAAGACCTGCCTGAAACCGAACTGCCCGCTGTTCTGCAGCCGGTCG
CGGAGGCCATGGATGCGATCGCTGCGGCCGATCTTAGCCAGACGAGCGGGTTCGGCCCATTCGG
ACCGCAAGGAATCGGTCAATACACTACATGGCGTGATTTCATATGCGCGATTGCTGATCCCCAT
GTGTATCACTGGCAAACTGTGATGGACGACACCGTCAGTGCGTCCGTCGCGCAGGCTCTCGATG
AGCTGATGCTTTGGGCCGAGGACTGCCCCGAAGTCCGGCACCTCGTGCACGCGGATTTCGGCTC
CAACAATGTCCTGACGGACAATGGCCGCATAACAGCGGTCATTGACTGGAGCGAGGCGATGTTC
GGGGATTCCCAATACGAGGTCGCCAACATCTTCTTCTGGAGGCCGTGGTTGGCTTGTATGGAGC
AGCAGACGCGCTACTTCGAGCGGAGGCATCCGGAGCTTGCAGGATCGCCGCGGCTCCGGGCGTA
TATGCTCCGCATTGGTCTTGACCAACTCTATCAGAGCTTGGTTGACGGCAATTTCGATGATGCA
GCTTGGGCGCAGGGTCGATGCGACGCAATCGTCCGATCCGGAGCCGGGACTGTCGGGCGTACAC
AAATCGCCCGCAGAAGCGCGGCCGTCTGGACCGATGGCTGTGTAGAAGTACTCGCCGATAGTGG
AAACCGACGCCCCAGCACTCGTCCGAGGGCAAAGGAATAGCCCGGGAAAA

Finally, we subcloned the fragments into #1068 vector (obtained from Dr. Alois Schweighofer, MFPL, Vienna, Austria) and used them for *Agrobacterium* mediated transformation of *Arabidopsis thaliana* plants by floral dip.
